# A unicellular relative of animals generates an epithelium-like cell layer by actomyosin-dependent cellularization

**DOI:** 10.1101/563726

**Authors:** Omaya Dudin, Andrej Ondracka, Xavier Grau-Bové, Arthur A. B. Haraldsen, Atsushi Toyoda, Hiroshi Suga, Jon Bråte, Iñaki Ruiz-Trillo

## Abstract

In animals, cellularization of a coenocyte is a specialized form of cytokinesis that results in the formation of a polarized epithelium during early embryonic development. It is characterized by coordinated assembly of an actomyosin network, which drives inward membrane invaginations. However, whether coordinated cellularization driven by membrane invagination exists outside animals is not known. To that end, we investigate cellularization in the ichthyosporean *Sphaeroforma arctica*, a close unicellular relative of animals. We show that the process of cellularization involves coordinated inward plasma membrane invaginations dependent on an actomyosin network, and reveal the temporal order of its assembly. This leads to the formation of a polarized layer of cells resembling an epithelium. We show that this epithelium-like stage is associated with tightly regulated transcriptional activation of genes involved in cell adhesion. Hereby we demonstrate the presence of a selforganized, clonally-generated, polarized layer of cells in a unicellular relative of animals.

## Introduction

Cellularization of a coenocyte — a multinucleate cell that forms through sequential nuclear divisions without accompanying cytokinesis — is a specialized form of coordinated cytokinesis that results in cleavage into individual cells. Cellularization commonly occurs during development of animals and plants. In both, cellularization results in the formation of a multicellular, spatially organized epithelium-like structure. Despite the similarities, distinct mechanisms are involved in the cellularization of animal and plant coenocytes. During endosperm development in most flowering plants, coenocytes cellularize through cell wall formation around individual nuclei, forming a peripheral layer of cells surrounding a central vacuole (Hehenberger et al., 2012). This is coordinated by the radial microtubule system and is dependent on several microtubule-associated proteins (Pignocchi et al., 2009; Sorensen, 2002). In contrast, in a model animal coenocyte, the syncytial blastoderm of the fruit fly *Drosophila melanogaster*, cellularization is accomplished through plasma membrane invaginations around equally spaced, cortically positioned nuclei (Farrell and O’Farrell, 2014; Mazumdar and Mazumdar, 2002). This process is driven by a contractile actomyosin network (Mazumdar and Mazumdar, 2002). It depends on several actin nucleators, such as the Arp2/3 complex (Stevenson et al., 2002) and formins (Afshar et al., 2000). It also requires multiple actin-binding proteins, including myosin II (Royou et al., 2002), which mediates actin cross-linking and contractility, as well as septins (Adam et al., 2000; Cooper and Kiehart, 1996), cofilin (Gunsalus et al., 1995) and profilin (Giansanti et al., 1998). In addition, it depends on cell-cell adhesion proteins including cadherin, and alpha- and beta-catenin (Hunter and Wieschaus, 2000; Wang et al., 2004). This coordinated cellularization results in the formation of a single layer of polarized epithelial tissue, also known as cellular blastoderm (Mazumdar and Mazumdar, 2002). Such cellularization is common in insects, however, whether this mechanism of cellularization is found outside animals, remains unknown.

Among holozoans — a clade that includes animals and their closest unicellular relatives (Figure 1A) — ichthyosporeans are the only known lineage besides animals that forms coenocytes during their life cycles (Mendoza et al., 2002; de Mendoza et al., 2015). All characterized ichthyosporeans proliferate through rounds of nuclear divisions within a cell-walled coenocyte, followed by release of newborn cells (Marshall and Berbee, 2011, 2013; Suga and Ruiz-Trillo, 2013; Whisler, 1968). We have previously suggested that they undergo cellularization (Ondracka et al., 2018; Suga and Ruiz-Trillo, 2013). However, at present nothing is known about the ichthyosporean cellularization, and whether it involves animal-like mechanisms.

**Figure 1.**
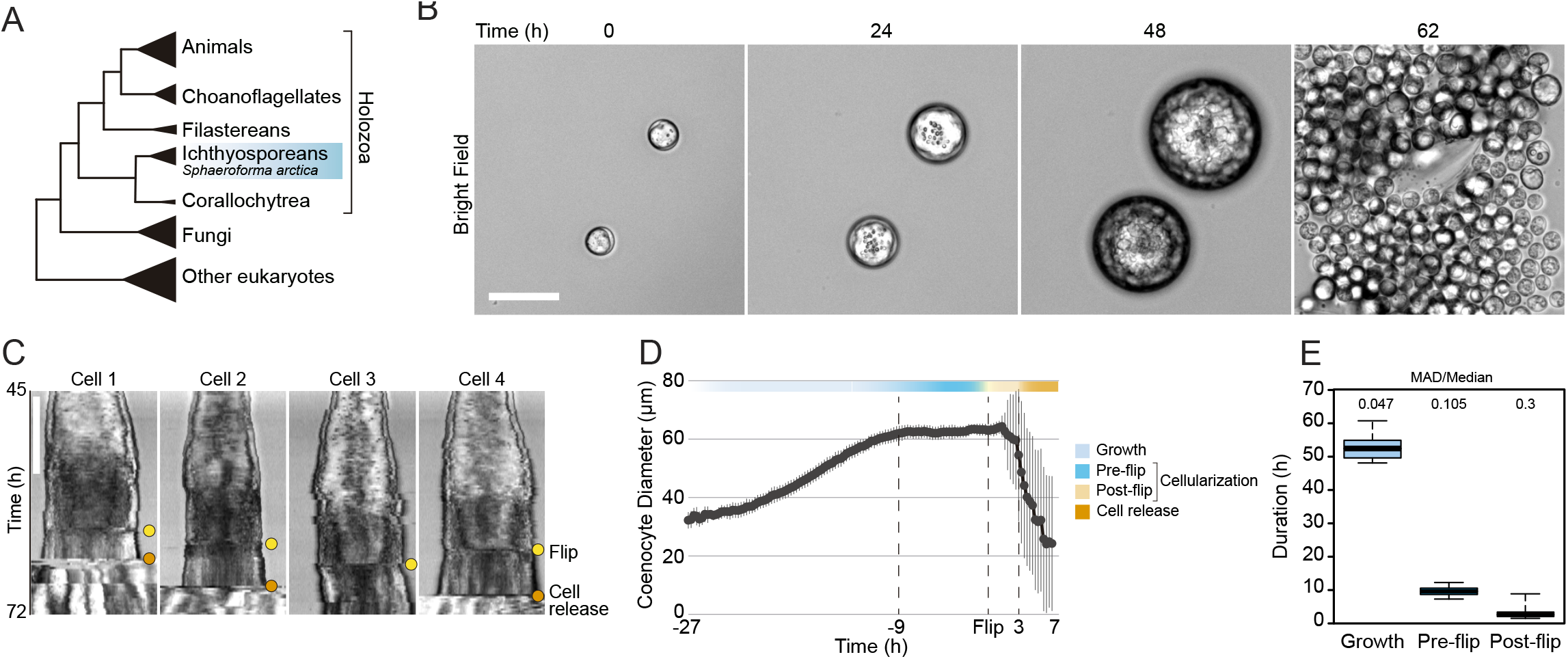
Cellularization dynamics in *Sphaeroforma arctica*. (A) Phylogenetic position of the ichthyosporean *Sphaeroforma arctica* in the tree of life (B) Time-lapse images of the life-cycle of *S. arctica* show cell-size increase prior to cellularization and release of new-born cells. Associated with Movie S2. Bar, 50μm. (C) Kymographs of 4 distinct cells undergoing cellularization with the time of flip (yellow) and cell release (orange) indicated for each. Bar, 50μm. (D) Cell diameter over time of 65 single cell traces aligned to Flip reveals distinct cell stages: Growth, Pre and post-flip and cell release. (E) Duration of growth, cellularization and post-flip represented as box-plots (N°cells > 100). MAD (Median absolute deviation) over median is used as a measure of variability.

Here, we comprehensively characterized cellularization in the ichthyosporean *Sphaeroforma arctica*, in which we have previously shown that coenocytic cycles can be synchronized (Ondracka et al., 2018). We used imaging, transcriptional profiling, and pharmacological inhibition, to study the gene expression dynamics, morphological rearrangements, and mechanisms of cellularization. We found that cellularization is accomplished by inward plasma membrane invaginations driven by an actomyosin network, forming a polarized layer of epithelium-like cells. Time-resolved transcriptomics revealed sharply regulated expression of cell polarity and cell adhesion genes during this epithelium-like stage. Finally, we show that this process depends on actin nucleators and Myosin II, and we reveal the temporal order of the actomyosin network assembly. Together, these findings establish that cellularization of ichthyosporeans proceeds by an animal-like mechanism.

## Results

### Growth and cellularization in *S. arctica* are temporally separated stages of the coenocytic cycle

To determine the timing of cellularization in synchronized cultures, we established long-term live imaging conditions. Individual coenocytes exhibit growth in cell size until approximately 60 hours, after which they undergo release of newborn cells (Figure 1B, Movie S1). This was consistent with previous results in bulk cultures (Ondracka et al., 2018), confirming that our experimental setup for long-term live imaging faithfully reproduces culture growth. However, by measuring the diameter of the coenocytes, we observed that newborn cell release occurred with somewhat variable timing (Figure S1A, Movie S1).

Time-lapse imaging revealed that prior to cell release, the coenocytes darken along the periphery, and the dark front begins to advance towards the center (Movie S2). Afterwards, we observed an abrupt internal morphological change in the coenocyte, when the front disappears. We termed this event “flip” (Movie S2). The flip occurred in all the coenocytes and can be reliably detected on kymographs (Figure 1C). Aligning individual coenocyte size traces to this specific common temporal marker, we observe that coenocytes stop growing approximately 9 hours before the flip (Figure 1D and 1E). This shows that the growth stage and cellularization are temporally separated. The cellularization can be divided into a temporally less variable pre-flip phase (~9 hours) and a variable post-flip phase (Figure 1E and S1C). Taken together, these results show that growth and cellularization form temporally distinct stages of the coenocytic cycle of *S. arctica*. This provides a temporal framework in which to characterize phenotypically distinct stages of cellularization.

### The cortical actin network establishes sites of membrane invagination and generates an epithelium-like layer of cells

To assess whether cellularization in *S. arctica* involves encapsulation of nuclei by plasma membranes, we imaged the plasma membrane using live time-lapse imaging in the presence of the lipid dye FM4-64 (Betz et al., 1996). We observed a rapid increase in FM4-64 intensity at the periphery of the coenocyte 30 minutes prior to flip (Figure 2A, panel II, Movie S3 and S4). The plasma membrane invagination sites formed at the periphery and progressed synchronously and rapidly from the outside toward the center of the coenocyte, forming polarized, polyhedral cells (Figure 2A, panels II-V, Movie S3 and S4). Finally, following flip, cells lost their polyhedral shape and became round, suggesting that they were no longer attached to each other (Figure 2A, panel VI, Movie S3 and S4).

**Figure 2.**
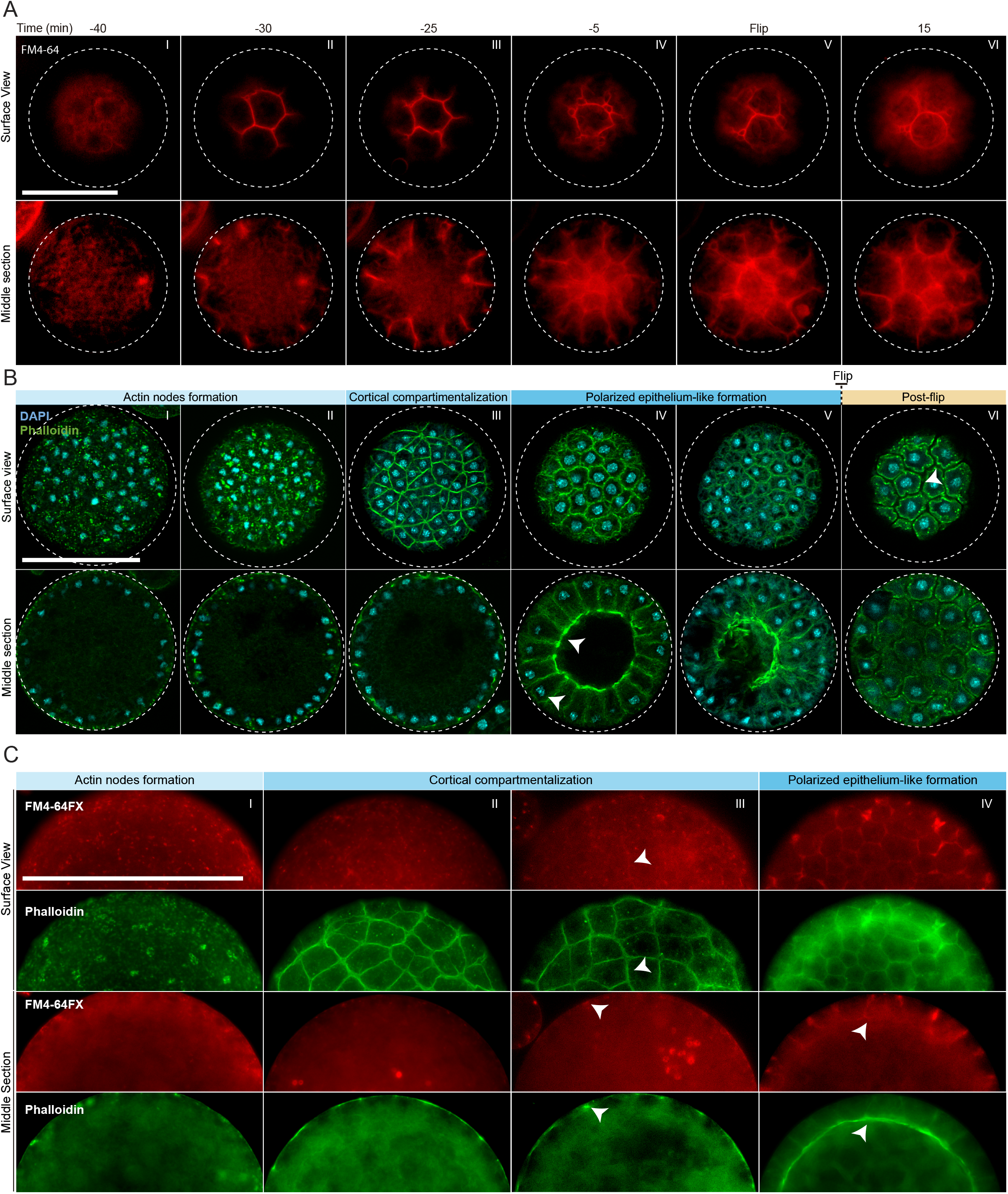
Actin cytoskeleton and plasma membrane dynamics during cellularization of *S. arctica*. (A) Dynamics of plasma membrane invaginations during cellularization. Live-cells, pre-grown for 58 hours, were stained with FM4-64 (10μM) and imagedusing epifluorescent microscopy with a 5 minutes interval. Bar, 50μm. (B) Spatio-temporal organization of the actin cytoskeleton, nuclei and cells during cellularization. Synchronized cells of *S. arctica*, pre-grown for 48 hours, were fixed every hour for 14 hours and stained with phalloidin and DAPI to reveal cytoskeletal dynamics during cellularization. All cells were imaged using confocal microscopy.In panel IV, arrows indicate higher actin signal intensity on the internal side and that nuclei are localized close to the cortex indicating that the epithelium-like layer of cells is polarized. In panel VI, an arrow indicate that cells are apart from each other after flip. Bar, 50μm. (C) Actin network is established prior to plasma membrane invaginations. Synchronized cells of *S. arctica*, pre-grown for 54 hours, were fixed every 2 hours for 10 hours and stained with both the membrane dye FM4-64FX and phalloidin. Arrows show sites of colocalization between both markers at the onset of plasma membrane invaginations. Bar, 50μm.

In animal coenocytes, plasma membrane invagination is associated with dynamic organization of the actomyosin cytoskeleton (Mazumdar and Mazumdar, 2002). To investigate actin dynamics during cellularization, we took advantage of the timeline described above and imaged coenocytes that were fixed and stained for actin and nuclei (using phaloidin and DAPI, respectively) at different time points during cellularization (Figure 2B, S2A and S2B). During growth, actin localized exclusively as small patches at the surface of the coenocyte (Figure S2A, panel I). Only at the onset of cellularization, multiple actin patches increased in size to form actin nodes (Figure 2B, panel I and II, and S2A, panel II). This phase was followed by cortical compartmentalization surrounding the nuclei through gradual formation of an actin filament network solely at the cortex of the coenocyte (Figure 2B, panel III, and S2A, panels III and IV). Following this cortical compartmentalization, an epithelium-like cell layer was formed by inward growth of the actin filaments from the cortex to generate a structure resembling the cellular blastoderm in *Drosophila* (Figure 2B, panels IV and S2A, panels V and VI). During this stage, the actin signal intensity was uneven and higher on the internal side (Movie S5), and nuclei were localized close to the cortex, indicating that cells in this epithelium-like layer are polarized (Figure 2B, panel IV, Movie S5). The epithelium-like layer progressively grew towards the center of the coenocyte to fill the cavity, until the center space was occupied (Figure S2A, panel VII). After flip, similar to the plasma membrane organization mentioned above, the epithelium-like layer of cells was reorganized to form spherical cells (Figure 2B, panel VI, and S2A, panel VIII).

To determine the order of actin organization and plasma membrane invaginations, we assessed the localization of both actin and plasma membrane in fixed samples using phalloidin and a fixable analog of FM4-64 (FM4-64FX). We found that the cortical actin network formed prior to the appearance of the membrane dye (Figure 2C, panel II).

The intensity of FM4-64FX labeling also increased and co-localized with the underlying actin network around the time of plasma membrane invagination (Figure 2C, panels III and IV). This suggests that the cortical actin network determines the site of plasma membrane invagination.

Finally, to determine the timing of cell wall formation, we stained the cells with calcofluor. We observed that labeling co-localized with the membrane marker FM4-64FX around individual cells in the post-flip stage (Figure S2C). This shows that the newborn cells already build the cell wall before the release, as was suggested previously in other *Sphaeroforma* species (Marshall and Berbee, 2013).

In early insect embryos, cellularization of the syncytial blastoderm occurs through actin-dependent invagination of the plasma membrane. Here, we demonstrate that the cellularization of the ichthyosporean coenocyte also involves active actin reorganization and membrane invagination at the site of actin cytoskeleton. Additionally, this results in the formation of a polarized epithelium-like layer of cells with an internal cavity that morphologically resembles the cellular blastoderm of insects.

### Cellularization is associated with extensive sequential transcriptional waves and is associated with evolutionarily younger transcripts

To gain insight into the regulation of the cellularization of *S. arctica*, we performed RNA-seq data on synchronized cultures with high temporal resolution, and comprehensively analyzed the dynamics of transcription, alternative splicing, and long intergenic non-coding RNAs (lincRNAs). Because the published genome assembly of *S. arctica* (Grau-Bové et al., 2017) was fragmented and likely resulted in incomplete gene models, we first re-sequenced the genome combining the Illumina technology with the PacBio long read sequencing technology. The final assembly sequences comprised 115,261,641 bp, and the metrics were greatly improved compared to the previous assembly (Grau-Bové et al., 2017) (Table S1). *Ab initio* gene annotation resulted in the discovery of novel ORFs due to the absence of repetitive regions in the previous assembly. RNA-seq data was used to improve the ORF prediction, to define the 5’ and 3’ untranslated regions, and to discover lincNAs. In total, 33,682 protein coding genes and 1,071 lincRNAs were predicted using this combined pipeline (see methods). This final transcriptome assembly was used as the reference transcriptome for further analysis.

To perform the time-resolved transcriptomics, we isolated and sequenced the mRNA from two independent synchronized cultures at 6-hour time intervals during the entire coenocytic cycle, encompassing time points from early 4-nuclei stage throughout growth and cellularization stages until the release of newborn cells (Figure S3A).

We first analyzed transcript abundance during the time series. The majority of the transcriptome (20,196 out of 34,753 predicted protein-coding and lincRNA genes) was transcribed at very low levels (mean expression throughout the time course <0.5 tpm [transcripts per million]) and were removed for the subsequent clustering analysis. Clustering of transcript abundance data from both biological replicates revealed a clear separation between the transcriptomes of the growth stage (12h to 42h time points) and the cellularization stage (48h to 66h time points) (Figure 3A). Furthermore, we observed that the transcriptome patterns between replicates 1 and 2 were shifted by 6 hours from 48 hours onwards (Figure 3A), presumably due to differences in temperature and conditions influencing the kinetics of the coenocytic cycle. We thus adjusted the time of the second replicate by 6 hours according to the clustering results, although we emphasize that the clustering analysis did not depend on time. Among the expressed transcripts (defined as mean expression levels higher than 0.5 tpm across all samples; in total 13,542 coding genes and 1,015 lincRNAs), consensus clustering using Clust (Abu-Jamous and Kelly, 2018) extracted 9 clusters of co-expressed transcripts with a total of 4,441 protein coding genes (Figure 3A), while the rest of the transcripts were not assigned to any coexpression cluster. The assigned cluster membership was robust to using either of the replicates or averaging (Figure S3B). Visualization by heatmap and t-SNE plot separated the clusters into two meta-clusters containing the growth stage (clusters 1-3, totaling 2,197 genes) and cellularization stage (clusters 4-9, totaling 2,314 genes) clusters (Figure 3C and 3D). Among the cellularization clusters, we see a clear temporal separation of different transcriptional waves, with clusters 5, 6 and 7 representing the three largest clusters of co-expressed transcripts of early, mid and late cellularization, respectively (Figure 3B and 3D). Altogether, this shows that cellularization is associated with extensive sharp transcriptional activation in multiple temporal waves, in total affecting 17% of the expressed transcriptome.

**Figure 3.**
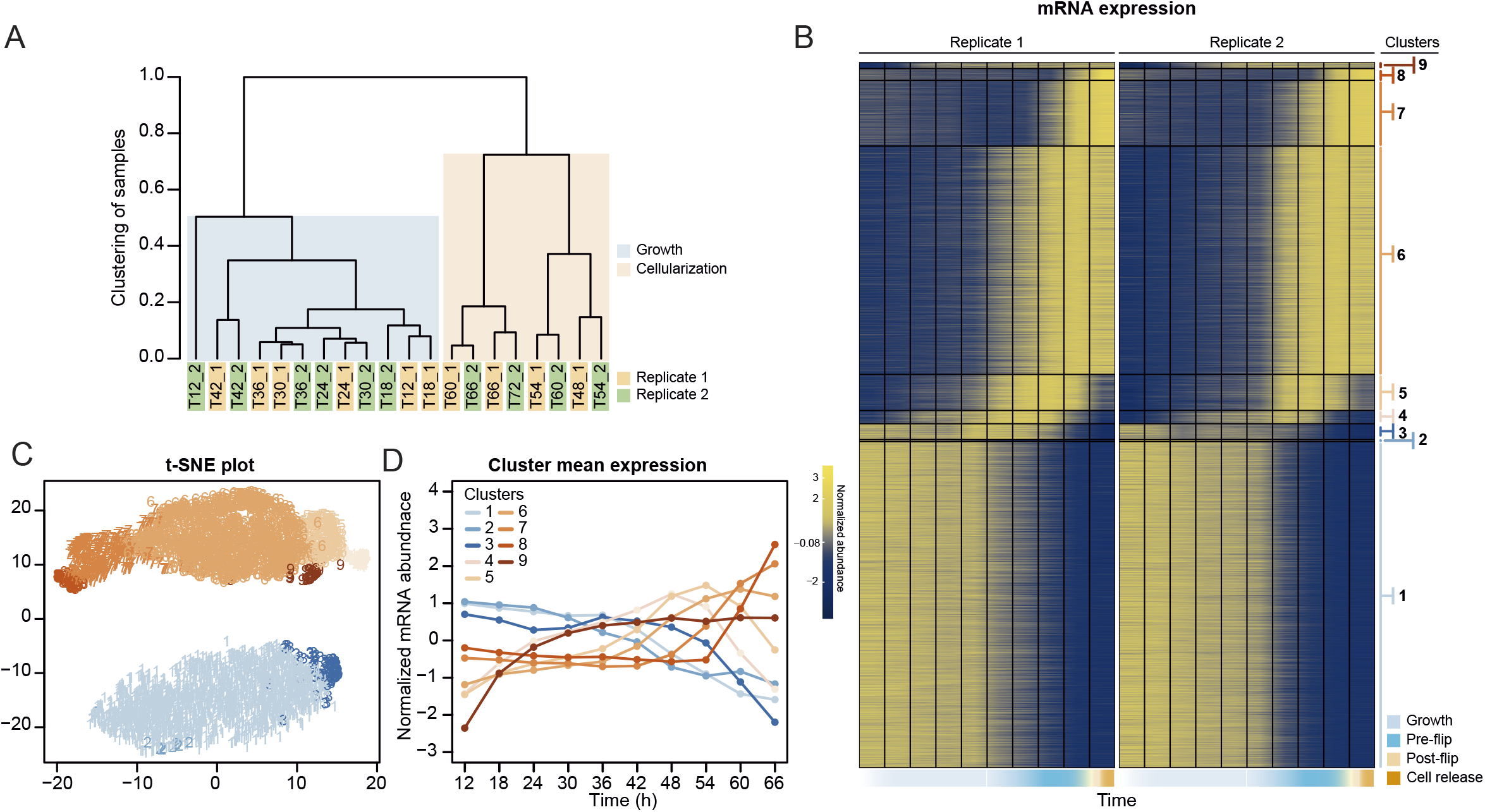
Transcriptional dynamics across the *S. arctica* life cycle. (A) Hierarchical clustering of time point samples by Euclidian distance of spearman correlation coefficient. The sample T48_2 is missing due to technical reason. (B) A heatmap of 4,441 coding genes that were clustered into 9 clusters. (C) A t-SNE plot of clustered genes. (D) Mean expression profile of each gene expression cluster.

In parallel to transcript abundance dynamics, we also assessed alternative splicing (AS) across the time series. This analysis identified 2,022 genes affected by intron retention (12.9% of all intron-bearing genes, totaling 4,310 introns), 914 by exon skipping (12.3% of genes with >2 exons, 1,206 exons) and 44 with mutually exclusive exons (0.7% of genes with >3 exons, involving 118 exon pairs) in all samples (Figure S3C and S3D). Overall, neither the number of AS events nor the number of genes affected vary dramatically along the *S. arctica* growth cycle (Figure S3E-S3J). Analysis of AS events over time did not yield any discernible global dynamics, although we found a small number of events differentially present between the growth and cellularization stages (Figure S3K).

Interestingly, skipped exons were more likely to be in-frame (38.63%, compared to 30.33% of in-frame exons in genes with >2 exons, *p* = 4.34e-05, Fisher’s exact test) and yield non-truncated transcripts, a phenomenon commonly observed in animal transcriptomes but not in transcriptomes of other unicellular eukaryotes (Grau-bové et al., 2018). The transcripts affected by such in-frame exon skipping events are enriched in biological processes such as regulation of multicellular organismal processes, assembly of the focal adhesion complex and cell growth (Figure S3L). Overall, although pervasive, alternative splicing likely does not play a major role in regulation of the coenocytic cycle and cellularization of *S. arctica*.

Additionally, we analyzed the dynamics of lincRNA expression. Overall, lincRNAs represent ~3% of total transcript abundance across the time series (Figure S4A). Among the long non-coding RNAs, 70 lincRNA transcripts clustered with coding genes into temporally co-expressed clusters (Figure 4A). Sequence homology search revealed that 24 of the *S. arctica* lncRNAs were conserved in distantly related ichthyosporean species such as *Creolimax fragrantissima, Pirum gemmata* and *Abeoforma whisleri* (estimated to have diverged ~500 million years ago (Parfrey et al., 2011). This is a remarkable depth of conservation, since animal lincRNAs are not conserved between animal phyla (Bråte et al., 2015; Gaiti et al., 2015; Hezroni et al., 2015). Other lincRNAs were either specific to *S. arctica* (511) or conserved only in closely related *Sphaeroforma* species (536). Comparison of lincRNA by degree of conservation showed no notable difference in %GC content (Figure S4B) or dynamics of expression during the coenocytic cycle (Figure S4C). However, we found that conserved lincRNAs were on average longer than non-conserved ones (Figure S4D) and, strikingly, expressed at much higher levels (Figure 4B). Among the 24 deeply conserved lincRNAs, 5 clustered in the temporally coexpressed clusters, which is higher than expected by chance (p = 0.0166, Fisher’s exact test). Among these, 3 belonged to the cellularization clusters, including, lincRNA asmbl_31839, which has a remarkably high sequence similarity with its homologs from other ichthyosporeans (Figure S4E). Furthermore, its transcriptional regulation is independent of the transcriptional regulation of its neighboring coding genes (located within 3kb) (Figure 4C). In summary, we discovered deeply conserved lincRNAs that are expressed at high levels and are transcriptionally activated during cellularization.

**Figure 4.**
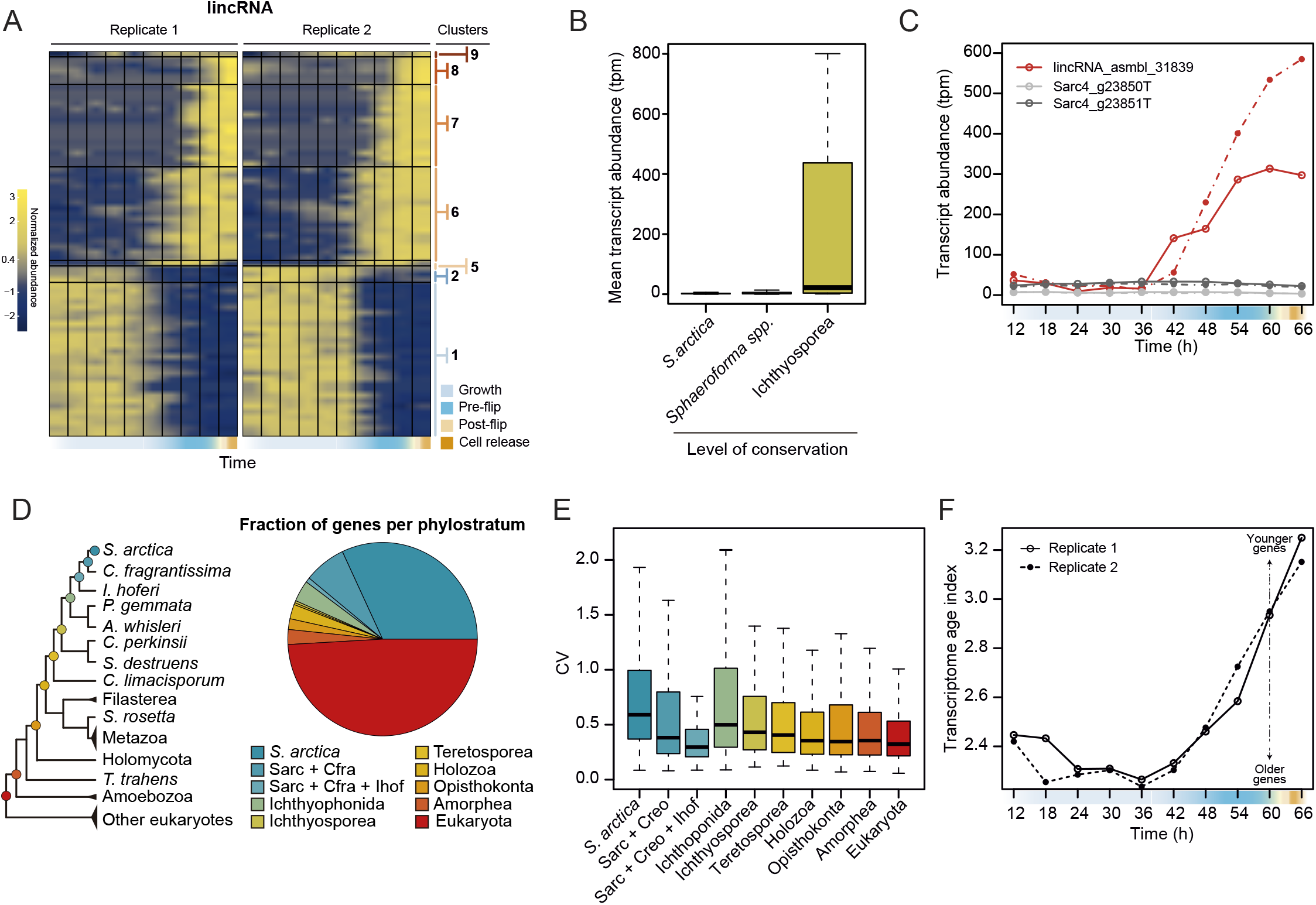
Dynamics of lincRNAs, alternative splicing, and gene phylostrata. (A) A heatmap of 70 long intergenic non-coding RNAs (lincRNAs) that co-cluster with coding genes. (B) mean expression level of the lincRNAs, binned by degree of conservation. (C) Expression of the conserved lincRNA, lincRNA: asmbl_31839 and the two coding genes located immediately upstream and downstream of it. (D) A phylogenetic tree indicating the 10 defined phylostrata, and a pie chart of fractions of all expressed genes per phylostratum (E) Coefficient of variance of gene expression across the *S. arctica* life cycle, binned by phylostratum (F) Transcriptome age index for the *S. arctica* life cycle

Finally, to assess the evolutionary origins of the co-expression clusters, we used a phylostratigraphic analysis to classify genes into evolutionary gene age groups. We carried out orthology analysis of the predicted *S. arctica* proteome along with 30 representative species from the eukaryotic tree of life to identify “orthogroups” (i.e. groups of putative orthologs between species). *S. arctica* protein-coding genes clustered in 6,149 orthogroups representing 12,527 genes; the rest of the genes did not have an ortholog outside *S. arctica*.

Next, we inferred the age of each gene using Dollo parsimony (Csuros, 2010) to classify them into phylostrata (sets of genes with the same phylogenetic origin) (Figure 4D). Analysis of gene expression by phylostrata revealed a trend toward more variable expression throughout the coenocytic cycle in younger genes (Figure 4E), although their mean expression levels were lower (Figure S5A). Such a correlation has been observed in animal development, where developmentally regulated genes tend to be of younger origin (Domazet-Lošo and Tautz, 2010). Analysis of enrichment of gene phylostrata in each gene expression cluster (Figure S5B) showed that the growth clusters are enriched for paneukaryotic genes. In contrast, we find that the cellularization clusters were enriched for younger genes (Figure S5B and S5C). Importantly, we found that genes with ichthyosporean origins were significantly enriched in all three of the largest cellularization clusters (Figure S5B and S5C). Computing the transcriptome age index (Domazet-Lošo and Tautz, 2010) revealed an hourglass pattern, with older genes expressed at later stages of growth, and younger genes expressed during early growth and cellularization (Figure 4F). Such an hourglass transcriptomic pattern has previously been observed in animal development, where it reflects the morphological similarities and differences of embryos of different taxa (Domazet-Lošo and Tautz, 2010), and it has been suggested as a conserved logic of embryogenesis across kingdoms (Quint et al., 2012). Despite this, we currently do not have a morphological explanation for this transcriptional hourglass pattern in ichthyosporeans. Altogether, the phylostratigraphic analysis suggests that cellularization is a comparatively younger process, whereas the growth stage represent an evolutionarily ancient process.

### Temporal co-expression of actin cytoskeleton, cell adhesion and cell polarity pathways during cellularization

To functionally assess the gene expression clusters, we also carried out gene ontology (GO) enrichment analysis (Table S2). The largest growth cluster (cluster 1) was enriched in GO terms related to cell growth and biosynthesis. Early and mid-cellularization clusters 5 and 6 were enriched for GO terms related to membrane organization and actin cytoskeleton. Late cellularization clusters were, in addition to GO terms related to actin, also enriched for GO terms related to cell adhesion and polarity. Given that these processes play a major role during cellularization of the insect blastoderm, we investigated the expression pattern of homologs of known regulators of cellularization in *Drosophila*.

In the *Drosophila* blastoderm, cellularization depends on the spatial organization of both the microtubule and actin cytoskeleton. It involves several microtubule and actin binding proteins, including kinesins, Myosin II, Myosin V, profilin (Chickadee), cofilin (Twinstar) and Septins, and the conserved family of Rho GTPases (Mazumdar and Mazumdar, 2002).

In *S. arctica*, we found that the expression of tubulins and kinesins, except for one (*S. arctica* Kinesin 2), is constant throughout the coenocytic cycle (Figure 5A and 5B). On the other hand, we found many actin-associated genes dynamically expressed. All actin nucleators of the formin family and members of the Arp2/3 complex peaked during cellularization (Figure 5C). In contrast to formins, which largely exhibited sharp peaks, the gene expression of the Arp2/3 complex was initiated earlier and increased gradually (Figure 5C). Septins, Cofilin, Profilin and Myosin V were temporally co-expressed during mid-cellularization and peaked at the same time as actin nucleators (Figure 5D), whereas myosin II, which has a role in organizing actin filaments and contractility, peaked later (Figure 5D). We likewise found the expression of all four members of the Cdc42/Rho1 orthogroup to be sharply activated during late cellularization (Figure 5E). Temporal co-expression of these genes suggests that the cellular pathways responsible for organizing the cytoskeleton and cell polarity in *Drosophila* cellularization are also involved in the cellularization process of *S. arctica*.

**Figure 5.**
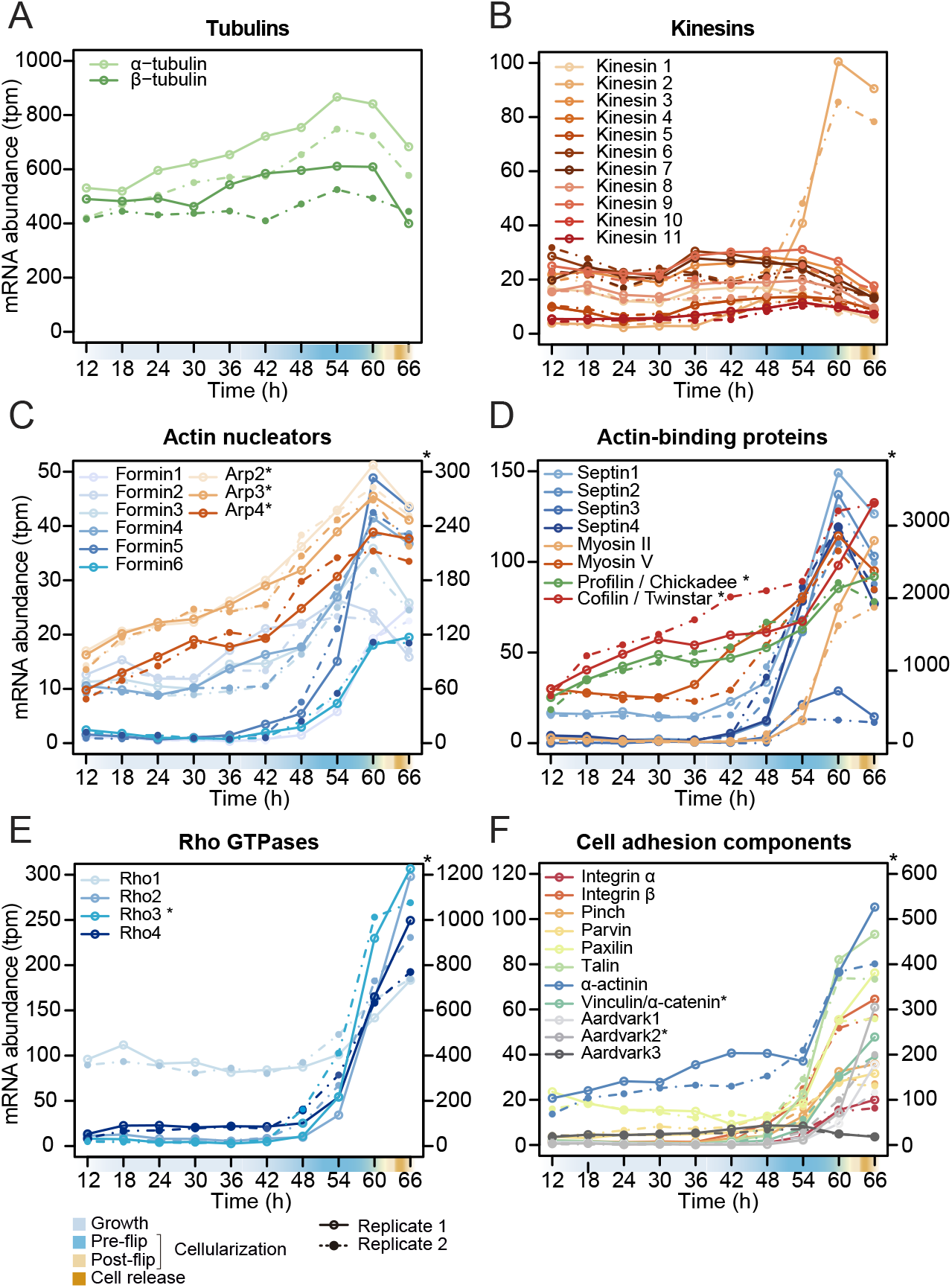
Temporal transcript abundance of cytoskeletal, cell polarity and cell adhesion genes. (A-F) Gene expression of indicated genes across the *S. arctica* life cycle

In addition to cytoskeletal organization, formation of a polarized epithelium also requires cell adhesion (Rodriguez-Boulan and Macara, 2014). Integrins and the cytoplasmic members of the integrin adhesome mediate cell-matrix adhesion in animal tissues. We observed that in *S. arctica*, both integrin receptors, alpha and beta, as well as all cytoplasmic members, are temporally co-expressed during late cellularization (Figure 5F). Beta and alpha catenin, together with cadherins in animals, mediate cell-cell adhesion in epithelial tissues (Rodriguez-Boulan and Macara, 2014). In *S. arctica*, we found three copies of Aardvark, a homolog of beta-catenin (Murray and Zaidel-Bar, 2014), as well as one homolog of alpha-catenin (Miller et al., 2013). We found that the expression of two out of three Aardvark transcripts, as well as the expression of the alpha catenin homolog, peaked during late cellularization (Figure 5F). This suggests that both cell-matrix and cell-cell adhesion pathways play a role in the establishment of the polarized epithelium-like structure during the late cellularization.

### The actomyosin cytoskeleton is essential for cellularization

Finally, we tested whether disrupting the cytoskeleton would lead to defects in cellularization. In the absence of genetic tools, we used small inhibitory molecules that target specific conserved cytoskeletal components. We synchronized the cultures and added the inhibitors at the onset of cellularization (54 h time point). We first assessed the role of the microtubule cytoskeleton during cellularization by adding carbendazim (MBC), a microtubule depolymerizing agent (Castagnetti et al., 2007). Microtubule inhibition did not prevent plasma membrane invagination, although it resulted in a delay of flip and release of newborn cells (Figure 6A and 6B, and S6A, Movie S6 and Movie S7).

**Figure 6.**
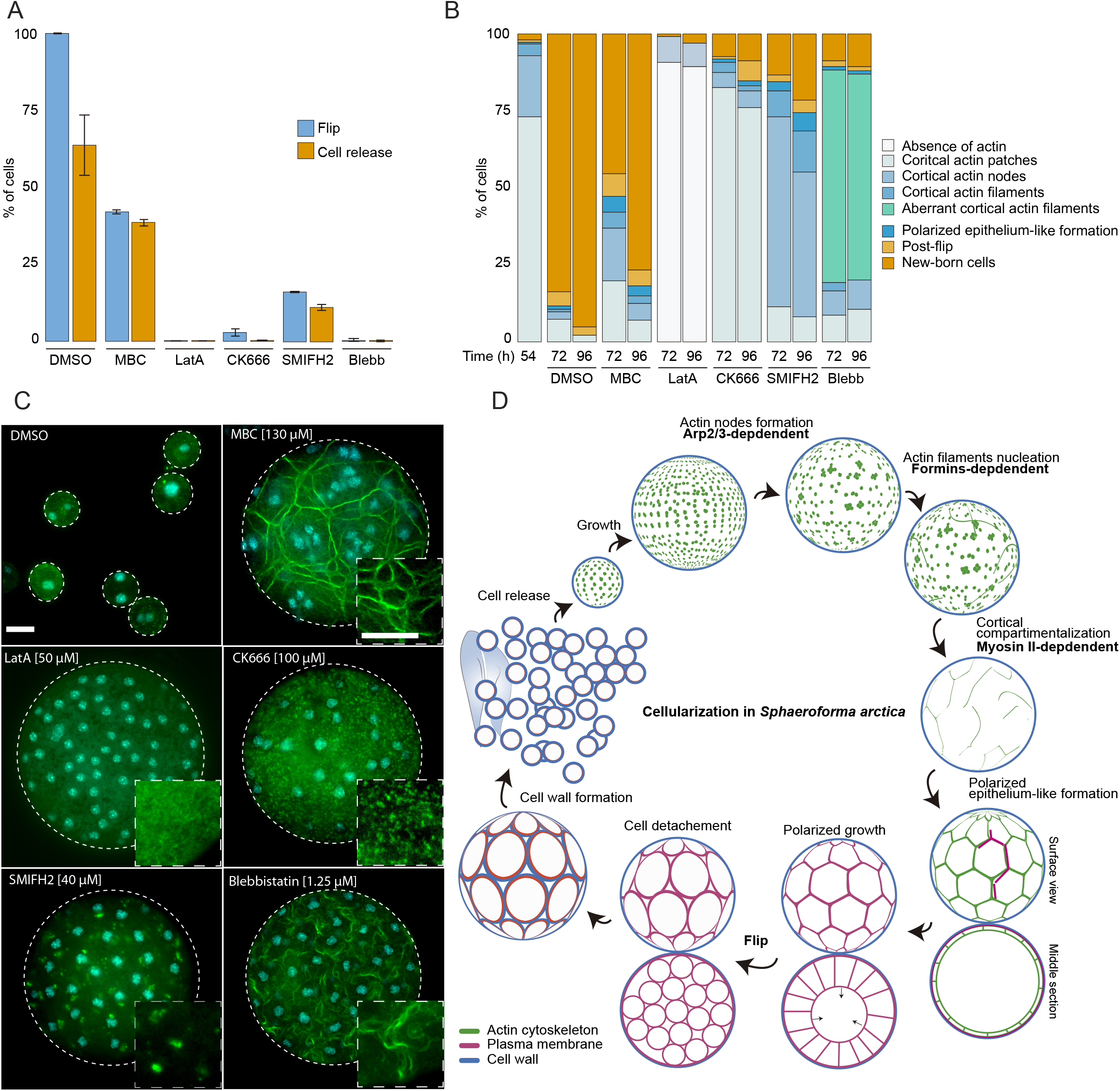
The actomyosin network organization is crucial for cellularization of *S. arctica*. (A) Depolymerization of microtubules and actin, as well as inhibition of Arp2/3, formins and myosin II affect “flip” and release of new born cells. Synchronized cells of *S. arctica*, pre-grown for 54 hours, were imaged for 24 hours in presence of multiple pharmacological inhibitors (DMSO as a control, MBC, Latrunculin A, CK666, SMIFH2, Blebbistatin). Cells undergoing flip and cell release throughout the duration of the experiment were measured (N>400 cells, Error bars are standard deviation from three independent experiments). (B) Temporal functions of the Arp2/3 complex, formins and myosin II in distinct stages of cellularization. Synchronized cells of *S. arctica*, pre-grown for 54 hours were subject to multiple pharmacological inhibitors treatments (DMSO as a control, MBC, Latrunculin A, CK666, SMIFH2, Blebbistatin) and fixed and stained with phalloidin and DAPI after 24h and 48 hours of treatments. Phalloidin staining allows us to measure the fraction of cells exhibiting different actin and cellular structures throughout cellularization. (C) The different actin structures observed when cells are treated with multiple pharmacological inhibitors treatments (DMSO as a control, MBC, Latrunculin A, CK666, SMIFH2, Blebbistatin). Bar, 10μm. (D) A model representing the actin cytoskeleton, plasma membrane and cell wall at different stages of the cellularization process in *S. arctica*, indicating sequential steps of actin remodeling mediated by Arp2/3, formins and Myosin II.

Furthermore, by staining the MBC-treated cells with DAPI and phalloidin, we observed a loss of the uniform spacing of the nuclei and actin filaments during the cortical compartmentalization stage of cellularization (Figure 6C). After MBC treatment, newborn cells varied in size and number of nuclei (Figure S6B, and S6C, Movie S6). These results suggest that the microtubule cytoskeleton is not essential for plasma membrane invagination but is crucial for nucleus and actin filament positioning at the cortex of the coenocyte.

Next, we sought to disrupt actin polymerization using either the broad actin depolymerizing agent Latrunculin A (LatA) (Braet et al., 1996), the Arp2/3 inhibitor CK666 (Hetrick et al., 2013), or the formin inhibitor SMIFH2 (Kim et al., 2015b). Cells treated with Latrunculin A lacked any actin patches or actin filaments and failed to undergo flip or produce newborn cells (Figure 6A-6C, Figure S6A, Movie S6 and S7). Furthermore, plasma membrane invagination did not occur after LatA treatment (Movie S6 and S7). In contrast, CK666-treated cells formed cortical actin patches, but were not able to form actin nodes, and they were unable to generate plasma membrane invaginations (Figure 6A-6C, Movie S6 and S7). These results show that the actin cytoskeleton and Arp2/3-mediated actin nucleation are required for the formation of actin nodes, the first step in the cellularization of *S. arctica*. In contrast, the formin inhibitor SMIFH2 did not block the formation of actin nodes, but it prevented the formation of actin filaments in the later stages (Figure 6A-6C). In addition, the plasma membrane invagination did not occur (Figure 6A-6C, Figure S6A, Movie S6 and S7), although we note that a small fraction of cells was not affected. This suggests that formins play a role in the nucleation of actin filaments after the formation of actin nodes.

Finally, we assessed the role of myosin II, using blebbistatin, an inhibitor of myosin II ATPase activity (Kovács et al., 2004). Blebbistatin treatment blocked plasma membrane invaginations and prevented cellularization (Figure 6A-6C, Figure S6A, Movie S6 and S7). Although blebbistatin-treated cells were able to form actin filaments, they had an aberrant wavy cortical actin filament network, suggesting that inhibition of myosin II causes loss of actin crosslinking and actin network contractility (Figure 6C). Taken together, these results indicate that the actomyosin apparatus is essential for cellularization in *S. arctica* and reveal a temporal sequence of involvement of Arp2/3 complex, formins and myosin II (Figure 6D). This temporal sequence is reflected in the relative timing of expression of Arp2/3, formins and myosin II genes (Figure 5C and 5D).

## Discussion

To address whether animal-like cellularization leading to formation of an epithelium exists outside animals, we performed imaging, transcriptomic analysis and pharmacological inhibition in a unicellular relative of animals, the ichthyosporean *Sphaeroforma arctica* (Figure 6D). We show that at the onset of cellularization, Arp2/3 complex mediates the formation of actin nodes at the cortex of the coenocyte. This is followed by formin-dependent nucleation of actin filaments. These actin filaments are then crosslinked in a myosin II-dependent manner, which results in formation of a cortical actomyosin network that surrounds evenly-spaced nuclei. This spatial organization of both the actomyosin network and nuclei depends on microtubules. The localization of the actomyosin network determines the sites of plasma membrane invaginations and drives their inward growth, resulting in the formation of a polarized epithelium-like layer of cells. This stage co-occurs with transcriptional activation of genes involved in cell-cell and cell-matrix adhesion. Polarized growth of the epithelium-like layer persists until the internal cavity is fully occupied, after which the polarized polyhedrally-shaped cells undergo flip, lose polarization and detach from each other. At this point, future newborn cells form the cell wall before they are released and initiate a new coenocytic cycle. Overall, our results show that the cellularization of *S. arctica* morphologically resembles the cellularization of the *Drosophila* blastoderm, a model for animal coenocytes. Furthermore, the roles of Arp2/3, formins and myosin II, which are all essential for plasma membrane invagination during the cellularization in *Drosophila* blastoderm, are conserved between ichthyosporean and animal coenocytes.

Unlike in *Drosophila* early embryos, our transcriptomic analysis suggests that transcriptional control plays an important role in regulation of the coenocytic cycle of *S. arctica*. Interestingly, we show that many genes expressed during cellularization of *S. arctica* emerged at the onset of ichthyosporeans. This strongly suggests that the general mechanisms of cellularization in *S. arctica* are likely conserved within ichthyosporeans. Altogether, our results argue that cellularization in ichthyosporeans requires both pathways conserved in animals, as well as pathways exclusive to ichthyosporeans. The later may include lincRNAs that we found to be remarkably conserved among ichthyosporeans and show high expression levels. Overall, our temporal gene expression dataset provides a good resource for further functional studies in *S. arctica* and other ichthyosporean species.

The epithelium is the first tissue that is established during embryogenesis and perhaps the first that emerged in evolution, thus representing the basic form of multicellular organization in animals. Epithelia are characterized by apical-basal polarity and cell-cell junctions, mediated by cadherins and catenins, that segregates the internal medium from the outside environment (Rodriguez-Boulan and Macara, 2014).

Epithelia were found in all animal lineages, including sponges (Leys et al., 2009). It has previously been suggested that the origin of epithelial structures predates the origin of animals, as a polarized epithelium-like structures, regulated by alpha- and beta-catenins, is also present in the slime mold *Dictyostelium discoideum* (Dickinson et al., 2011). Based on this finding, it has been suggested that all unikonts (Amorphea) — a clade including animals, its close holozoan unicellular relatives, as well as Fungi, Apusomonadida, Breviatea, and Amoebozoa — evolved from an ancestor with simple multicellular organization (Dickinson et al., 2012). However, epithelial structures have not yet been observed in other unikont lineages besides animals and slime molds (Brunet and King, 2017), raising questions whether the origin of these epithelial structures in these two relatively divergent lineages might have originated independently.

Here we argue that epithelium-like structures are generated during cellularization of ichthyosporeans. We show that the epithelium-like layer of cells is polarized due to the asymmetrical localization of cellular components and the direction of growth. Furthermore, despite lacking genetic tools to demonstrate the function of alpha and beta catenin, the sharp expression of catenins during the epithelium-like stage of cellularization strongly suggests a function in cell-cell junctions. However, whether this layer of cells functions to segregate the interior of the coenocyte from the outside environment remains to be addressed. Our results in ichthyosporeans lends support to the hypothesis that the origin of epithelial tissues predates the origin of animals. However, unlike the epithelium formation in social amoebae, which originates through aggregation (Bonner, 1998), ichthyosporean epithelium is generated clonally, such as epithelial tissues in all modern animals. Further studies to better understand the mechanisms of epithelium formation in ichthyosporeans will provide insight into the evolutionary origin of one of the defining features of animal multicellularity, the spatiotemporal cellular organization of tissues.

## Author contributions

O.D. and A.O. designed the study and set up the methodology. O.D. performed the microscopy and chemical inhibition experiments. A.O. performed the *S. arctica* RNA extraction, flow cytometry, and analyzed the gene expression data. A.T. and H.S. sequenced and assembled the *S. arctica* genome. X.G.-B. performed the genome annotation and alternative splicing analysis. A.A.B.H. and J.B. performed the de-novo transcriptome annotation, lincRNA prediction and conservation analysis. I.R.-T. provided supervision. O.D., A.O. and I.R.-T. wrote the original draft. All authors reviewed and edited the manuscript.

## Acknowledgments

We thank Sophie Martin, Aaron New, Daniel Richter and Sébastien Wielgoss for discussion and comments on the manuscript, Takaaki Kai for discussion on the ichthyosporean development, Eduard Ocaña for advice on the phylostratigraphic analysis, and Meritxell Antó for technical support. We also acknowledge the UPF Flow Cytometry Core Facility for assistance with flow cytometry, CRG Advanced light microscopy unit for support with confocal imaging, and the CRG Genomics Unit for mRNA library preparation and Illumina sequencing. This work is dedicated to the memory of Arthur Haraldsen, our dear friend and colleague, who tragically passed away during the final stages of the manuscript preparation.

This work was funded by European Research Council Consolidator Grant (ERC-2012-Co -616960) to I.R.-T.; MEXT KAKENHI (grant Nos. 221S0002 to A.T.; grants no 26891021 and 16K07468 to H.S.); and a Young Research Talents grant from the Research Council of Norway (grant no. 240284) to J.B.. O.D. was supported by a Swiss National Science Foundation Early PostDoc Mobility fellowship (P2LAP3_171815) and a Marie Sklodowska-Curie individual fellowship (MSCA-IF 746044). A.O. was supported by a Marie Sklodowska-Curie individual fellowship (MSCA-IF 747086).

## Methods

### Culture conditions

*Sphaeroforma arctica* cultures were grown and synchronized as described previously (Ondracka et al., 2018). Briefly, cultures were grown in Marine Broth (Difco BD, NJ, USA; 37.4g/L) in flasks at 12°C until saturation, producing small cells. These cells were then diluted into fresh media with low density (1:300 dilution of the saturated culture), resulting in a synchronously growing culture. Cultures of *S. gastrica, S. tapetis, S. nootkatensis, P. gemmata* and *A. whisleri* were similarly kept in Marine Broth (Difco BD, NJ, USA; 37.4g/L) at 12°C in dark conditions.

### *Sphaeroforma arctica* genome sequencing and assembly

Genomic DNA from *S. arctica* was extracted using QIAamp DNA Blood Midi Kit (Qiagen) from 300 mL culture incubated at 12 C° for 1 week in six 75 cm^2^ flasks. The Qubit (Invitrogen) quantification found ~150 μg genomic DNA in total.

A SMRTbell library for P6/C4 chemistry was constructed and was run on 32 SMRT cells in a PacBio RSII system (Pacific Biosciences), generating 2,209,004 subreads with a total of 25,164,714,269 bp. Raw subreads were first assembled using the FALCON_unzip assembler (v0.4.0) and the initial assembled sequences were polished by the Quiver integrated in SMRT analysis (v2.3.0). Genomic DNA was then sheared using a Focused-ultrasonicator (Covaris, Inc.). A paired-end library with an insert size of 600 bp was sequenced on an Illumina HiSeq2500 platform, producing 136,566,600 reads with read length of 250 bp. Paired-end reads were mapped against the polished sequences using the BWA mem (v0.7.12) followed by error-correction using Pilon (v1.22) (Walker et al., 2014).

### *Sphaeroforma arctica* gene annotation

We annotated predicted gene models in our scaffolds using BRAKER2 (Hoff et al., 2018). First we used STAR 2.7 (Dobin et al., 2012) to map all the RNA-seq samples to the genomic scaffolds, so as to obtain empirical evidence of gene bodies and guide the prediction of gene coordinates. These read mappings were then supplied to BRAKER2 in BAM format, which combied these external evidence with the gene coordinates predicted by Augustus 3.2.2 (Keller et al., 2011) (--noInFrameStop mode) and Genemark v4.21 (Lomsadze et al., 2005) to obtain a consolidated set of gene models in GFF format.

The transcriptome of *S. arctica* was assembled in order to add 5’ and 3’ UTR regions to the genome annotation and to search for long non-coding RNA-seq using data obtained here (see below).

The RNA-seq reads from the first time series replicate (excluding timepoint 30) were subjected to quality trimming and adaptor removal using Trimgalore (v0.4.5) (http://www.bioinformatics.babraham.ac.uk/projects/trimgalore/) and Trimmomatic v0.35 (Bolger et al., 2014). Trimgalore was first run to remove sequencing adaptors, allowing 0 mismatch and minimum adaptor length of 1 nt. Trimmomatic was run by trimming bases with phred score <20 from both ends. Furthermore, a sliding window of 4 bases was used to trim reads from the 5’ end when the mean phred score dropped below 20. Finally, the IlluminaClip option was used to search for remaining sequencing adaptors, allowing 2 seed mismatches, a palindrome clip threshold of 20 and minimum single match threshold of 7 nt. The quality filtered RNA reads were subsequently assembled using Trinity v2.5.1 (Grabherr et al., 2011), running both a *de novo* and a genome-guided assembly. To perform a genome-guided transcriptome assembly, we first mapped the RNA-seq reads against the genome using Hisat2 v2.1.0 (Kim et al., 2015a), in strand-specific mode with default parameters. Both Trinity assemblies were run in strand-specific mode while applying the jaccard clip option and otherwise default parameters. We next evaluated the strand-specificity of the assemblies by mapping RNA-seq reads back to the Trinity assemblies using scripts supplied within the Trinity software. Transcripts were subsequently removed if >80% of the reads mapped in the wrong direction. The PASA pipeline v2.3.1 (Haas et al., 2003) was then used to update the existing genome annotation. First, the assembled transcripts described above were mapped against the genome using the initial PASA script to make a temporary annotation file. This was performed with default parameters, using both Blat v35 (Kent, 2002) and Gmap v2015-09-29 (Wu and Watanabe, 2005) aligners, with transcripts specified as strand-specific. Lastly, the transcriptome-based annotations were compared with the existing genome annotation in a final PASA run, also using default parameters. In this step, the existing genes were expanded with UTR annotations and, in cases where a single RNA transcript covered multiple genes, these become merged into a single gene.

### RNA isolation, library preparation and sequencing

Synchronized cultures of *Sphaeroforma arctica* at 12°C were sampled every 6 hours for a total duration of 72 hours. Total RNA was extracted by Trizol and purified using the miRNeasy Mini Kit (QIAGEN) from ~50mL of culture at each time point. Libraries were prepared using the TruSeq Stranded mRNA Sample Prep kit. Paired-end 50bp read length sequencing was carried out at the CRG genomics core unit on an Illumina HiSeq v4 sequencer. We obtained between 19 and 32 M reads per sample. Transcripts were quantified using Kallisto software (Bray et al., 2016) with default parameters. To remove non-expressed genes, we filtered out transcripts that had a mean expression level of <0.5 tpm across all 20 samples. This resulted in a set of 14557 transcripts that were used for clustering.

For *S. gastrica* and *S. tapetis*, RNA purification was performed using the RNeasy kit (Qiagen), while for *S. nootkatensis, P. gemmata* and *A. whisleri*, RNA was purified using Trizol (Invitrogen, CA, USA). Strand-specific sequencing libraries were prepared using Illumina TrueSeq Stranded mRNA Sample Prep kit and sequenced on an Illumina HiSeq3000 machine (150 bp paired end).

### Clustering of the gene expression and gene ontology enrichment analysis in *S. arctica*

*Sphaeroforma arctica* transcripts were clustered by their gene expression profiles using Clust software (Abu-Jamous and Kelly, 2018) with default parameters and automatic normalization mode. This yielded 4511 transcripts (4441 coding genes and 70 lncRNA genes) that were clustered into 9 clusters, ranging from 41 to 2081 co-expressed genes. Gene expression profiles were visualized using superheat package in R (Barter and Yu, 2018). The tSNE plot was generated using Rtsne package.

Gene Ontology enrichments based on the GOs annotated with eggNOG mapper (Huerta-Cepas et al., 2017) were computed using the topGO R library (v. 2.34). Specifically, we computed the functional enrichments based on the counts of genes belonging to the group of interest relative to all annotated genes, using Fisher’s exact test and the elim algorithm for ontology weighting (Alexa et al., 2006).

### Transcriptome assembly of other unicellular holozoans

Raw reads (from sequencing libraries and SRA data) were processed with Trimmomatic (Bolger et al., 2014) to remove adapters and low-quality bases, by trimming bases with phred score <28 from both ends. Furthermore, a sliding window of 4 bases was used to trim reads from the 5’ end when the mean phred score dropped below 28. Finally, the IlluminaClip option was used to search for remaining sequencing adaptors, allowing 2 seed mismatches, a palindrome clip threshold of 28 and minimum single match threshold of 10 nt. All libraries were assembled denovo with Trinity (v2.3.2-2.5.1) (Grabherr et al., 2011) using default parameters. The assemblies of RNA-seq data were performed with the strand specific option, while assemblies based on SRA data were run in standard mode. For the *S. sirkka* assembly we also applied the jaccard clip option. Coding regions were predicted using TransDecoder v5.2.0 (Haas et al., 2013), by first extracting the longest possible ORFs (only top strand was searched in the strand-specific assemblies), on which likely coding region was predicted. Only the longest ORF was kept for each transcript.

### DNA isolation and genome assembly of *S. gastrica* and *S. tapetis*

Genomic DNA (gDNA) was isolated from *S. gastrica* and *S. tapetis*, by lysing cells on a FastPrep system (MP Biomedicals, CA, USA; 4m/s, 20s) followed by gDNA purification using the DNeasy kit (Qiagen, NRW, Germany) and subsequently sequenced on an Illumina HiSeq X system (150 bp paired end). Raw reads were subjected to quality trimming and adaptor removal as described above for RNA-seq data, with a phred score cut-off of 26.

The quality trimmed reads were subsequently error-corrected and assembled using Spades v3.10.0 (Nurk et al., 2013) applying kmer values 21, 33, 55, 77, 99 and 121, but otherwise default parameters. The resulting spades assemblies were scaffolded using L_rna_scaffolder (Xue et al., 2013) and polished with Pilon (Walker et al., 2014). L_RNA_Scaffolder was run by first mapping the respective transcriptome assemblies to the genome assemblies using Blat (Kent, 2002) which was inputted to L_rna_scaffolder. Next, we run Pilon by first mapping the quality trimmed genomic Illumina reads to the genome assembly using Bowtie2 (Langmead et al 2012). The resulting mapping file was then used in Pilon with default parameters. L_rna_scaffolder and Pilon were run repeatedly 5 times, followed by 3 final runs using only Pilon. These genome assemblies were used as reference genomes for genome-guided Trinity assemblies and PASA annotation as previously described for *S. arctica*.

### *S. arctica* long intergenic non-coding RNA identification and conservation analysis

Long intergenic non-coding RNAs (lincRNAs) were identified from the PASA annotation described above. First, transcripts shorter than 200 nt were discarded. Then, coding potential was evaluated using both TransDecoder (Haas et al., 2013) and CPC2 (Kang et al., 2017) with default parameters. All transcripts lacking coding potential were then compared with the Rfam database (Kalvari et al., 2017) to exclude other non-coding RNAs such as rRNAs and tRNAs. To exclude possible UTRs actually belonging to fragmented protein coding genes, a minimum genomic distance of 1000 bp from the closest protein coding gene was required. The remaining transcripts were compared with protein coding genes in the Swissprot database using Blastx (Altschul et al., 1990), and all sequences with a match better than e-value 1e-5 were removed.

Next, the potential lncRNA transcripts were blasted against the proteomes of *S. arctica* and other closely related ichthyosporeans (*Sphaeroforma tapetis, Sphaeroforma sirkka, Sphaeroforma nootkatensis, Sphaeroforma napiecek, Sphaeroforma gastrica, Creolimax fragrantissima, Pirum gemmata* and *Abeoforma whisleri*), and all transcripts with a match better than e-value 1e-5, were discarded. To remove transcripts resulting from potential spurious transcription, we required a minimum expression level of at least 1 tpm in at least one time series sample.

To search for possible conserved lncRNAs in other species, we performed Blastn searches against transcriptomes and genomes of several holozoan species (Richter et al., 2018) (Table S3). All genes providing hits with an e-value less than 1e-50 were considered to be a conserved homolog.

### Genome-wide analysis of alternative splicing

Each RNA-seq run was independently aligned to the *Sphaeroforma* genome using Hisat2 (Kim et al., 2015a), using default parameters except for longer anchor lengths to faciliate *de novo* transcriptome assembly (--dta flag).

The resulting alignments (in SAM format) were used to build sample-specific transcriptome assemblies with Stringtie2, using existing gene models (in GFF format, -G flag) as a reference, a minimum isoform abundance of 0.01 (−f flag) and a minimum isoform length of 50 bp (−m flag). For each sample, we only retained transcripts that overlapped with known genes in the final GFF file (using bedtools (Quinlan and Hall, 2010)). Then, we built a consolidated set of isoforms by pooling all sample-specific GFF annotations and the reference annotation using Stringtie2 (--merge flag), without imposing any limitation of minimum expression levels (−T flag set to 0), and retaining isoforms with retained introns (−i flag). We also calculated the expression levels at the isoform level using Salmon (Patro et al., 2017) (output in TPM). Then, we used SUPPPA2 (Trincado et al., 2018) to generate a set of alternative splicing events, for which their frequencies were calculated for each sample. Specifically, we used the consolidated set of isoforms (GFF format) to obtain a list of all possible AS events using SUPPA2 *generateEvents* mode (setting the output to exon skipping [SE], mutually exclusive exons [MX] and intron retention [RI]), and -l 10. Then, we used SUPPA psiPerEvent mode to calculate the PSI values of each AS event for each sample, using the expression levels of each isoform (obtained from Salmon) as a reference. Differential splicing was quantified by calculating the calculating the differential PSI values between the average of each sample group (growth stage [t=12h to t=48h] compared to cellularization stage [t=54h to t=66h]). p-values were obtained using the empirical significance calculation protocol described in SUPPA2.

After running SUPPA2, we produced functional annotations of the effect of each AS event on the final transcript. First, we annotated the protein domains in our consesus gene models using Pfam (version 31) and Pfamscan. These protein-level coordinates were converted to genomic coordinates using BLAT alignments (version 36, by aligning protein sequences to 6-frame translations of the genome) (Kent, 2002) and intersected with the genomic coordinates of AS events using the GenomicRanges and IRanges libraries (*findOverlapPairs* module) (Lawrence et al., 2013) from the R statistical framework (version 3.5.2). Second, we used the genomic coordinates of all AS events to recode each AS event as insertion/deletion variants that were amendable for analysis using the Variant Effect Predictor software (version 92.3) (McLaren et al., 2016) (SE were encoded as deletions, RI as insertions, and MX as insertion/deletion complex events). VEP produced a list of the effects of each AS variant (according to the Sequence Ontology nomenclature (Eilbeck et al., 2005)), using our consensus gene models as a reference.

### Phylostratigraphy analysis

First, we performed gene orthology assignment by searching for orthologs in a representative set of 30 publicly available eukaryotic proteome sequences (animals: *H. sapiens, S. kowalewskii, D. melanogaster, T. tribolium, N. vectensis, T. adherens, M. leidii and A. queenslandica;* choanoflagellates: *S. rosetta;* filastereans: *C. owczarczaki* and *M. vibrans;* teretosporeans: *S. arctica, C. fragrantissima, I. Hoferi, A. whisleri, C. perkinsii, P. gemmata, S. destruens, C. limatocisporum*, fungi: *F. alba, C. anguillulae, S. punctatus, M. verticillata* and *S. pombe;* other eukaryotes: *T. trahens, A. castellanii, D. discoideum, N. gruberi, T. thermophila, E. huxleyi, A. thaliana* and *P. yezoensis*) using orthofinder (Emms and Kelly, 2015) in Diamond mode. The generated orthogroups were used to determine orthologs of *S. arctica* genes in other species, unless a phylogeny of a specific gene family has been published. We then determine gene age of each orthogroup by Dollo parsimony implemented in Count software (Csuros, 2010) to classify the *S. arctica* proteome into 10 phylostrata, ranging from paneukaryotic to S. arctica-specific. The transcriptome age index was computed using a formula defined in (Domazet-Lošo and Tautz, 2010).

### Flow cytometry

Flow cytometry was performed as described previously (Ondracka et al., 2018) Cells were fixed in 4% formaldehyde, 1M sorbitol solution for 15 minutes at room temperature, washed once with marine PBS (PBS with 35 g/L NaCl), and stained with DAPI (final concentration 0.5 μg/mL) in marine PBS. Samples were analyzed using an LSRII flow cytometer (BD Biosciences, USA) and the data were collected with FACSDiva software. DAPI signal was measured using a 355nm laser with the 505nm longpass and 530/30nm bandpass filters. Approximately 2,000 events were recorded in each measurement. The flow cytometry data were processed and analyzed using FloJo software (Ashland, OR).

### Microscopy

Confocal microscopy of the spatiotemporal organization of actin in Figure 2A and Movie S5, was performed using a confocal laser scanning Leica TCS SP5 II microscope with an HC PL APO 63x/1.40 Oil CS2 oil objective. All remaining live and fixed images were obtained using a Zeiss Axio Observer Z.1 Epifluorescence inverted microscope equipped with Colibri LED illumination system and an Axiocam 503 mono camera. A Plan-Apochromat 63X/1.4 oil objective has been used for imaging fixed cells (Figures 2C, S2A, 6B, 6C), and an EC Plan-Neofluar 40x/0.75 air objective for supplementary Figure 2C, Movies S3 and S4 and an N-ACHROPLAN 20x/0.45na Ph2 air objective for live imaging in Figures 1C, 1B, S1B, Movie S1, S2, S6, S7.

### Cell fixation and staining

Cells were fixed using 4% formaldehyde and 250mM sorbitol for 30 minutes before being washed twice with PBS. For actin and nuclei staining phalloidin (Figures 2A, 2C, 6C, S2A, S2B), cells were span down at 1,000 rpm for 3 min after fixation and washed again three times with PBS before adding 10 μl of Alexa Fluor 488–phalloidin (Invitrogen) and DAPI at a final concentration of 5 μg/mL to 5 μl of concentrated sample. For plasma membrane and cell wall staining (Figure S2C), cells were incubated for 10 minutes with FM4-64FX (Invitrogen) at a final concentration of 10μM from 100× DMSO diluted stock solution and Calcofluor white (Sigma-Aldrich) at a final concentration of 5 μg/ml from a 200× stock solution prior to fixation. Cells were then fixed as previously mentioned and concentrated before being disposed between slide and coverslip.

For figures 5A, S5A and S5B, cells were pre-grown at 12°C for 48h prior to fixing every hour for a total of 14 hours.

### Live microscopy and pharmacological inhibitors

For live-cell imaging (Figures 1C-1F, 2B, 6A, S1A, S1B, S1C, Movies S1, S2, S4-S7) saturated culture was diluted 300x in fresh marine broth medium 1X inside a μ-Slide 4 or 8 well slide (Ibidi) at time zero. To ensure oxygenation during the whole period of the experiment, the cover has been removed. To maintain the temperature at 12°C we used a P-Lab Tek (Pecon GmbH) Heating/Cooling system connected to a Lauda Ecoline E100 circulating water bath. To reduce light toxicity, we used a 495nm Long Pass Filter (FGL495M-ThorLabs). Kymographs of cells undergoing cellularization in Figure 1D were constructed in ImageJ v1.46 by drawing a 3-pixel-wide (0.39 μm) line crossing the center of each cell.

For plasma membrane live staining (Figure 2B, Movie S4, S5), FM4-64 (Invitrogen) at a final concentration of 10μM from 100× DMSO diluted stock solution was added after 58 hours of growth. Treatment with pharmacological inhibitors was performed on 54 hours grown cells inside a μ-Slide 8 well slide (Ibidi) at 12 °C and lasted for 24 hours during which live microscopy was performed. Latrunculin A (Sigma-Aldrich) was used at final concentration of 50 μM from a stock of 20 mM in DMSO. CK666 (Sigma-Aldrich) was used at a final concentration of 100 μM from a stock of 10mM in DMSO. SMIFH2 (Sigma-Aldrich) was used at a final concentration of 40 μM from a stock of 10mM in DMSO. Blebbistatin (Sigma-Aldrich) was used at a final concentration of 1.25 μM from a stock of 2.5mM in DMSO. MBC (Sigma-Aldrich) was used at final concentration of 130 μg/ml from a stock of 2.5 mg/ml in dimethyl sulfoxide (DMSO).

### Image analysis

Image analysis was done using ImageJ software (version 1.52) (Schneider et al., 2012). For measurements of cell diameter in Figures 1E and S1A, we cropped movies to ensure having a single cell per movie. We then transformed the movies into binaries to ensure later segmentation. We then used particle analysis function in ImageJ with a circularity parameter set to 0.65-1 to quantify measure cell perimeter. As cells are spherical, we computed cell diameter as:

For quantification of fraction of cells in each stage of cellularization Figure S2B, we used the ObjectJ plugin in ImageJ (National Institutes of Health).

All Figures were assembled with Illustrator CC 2017 (Adobe).

---

**Figure S1.**
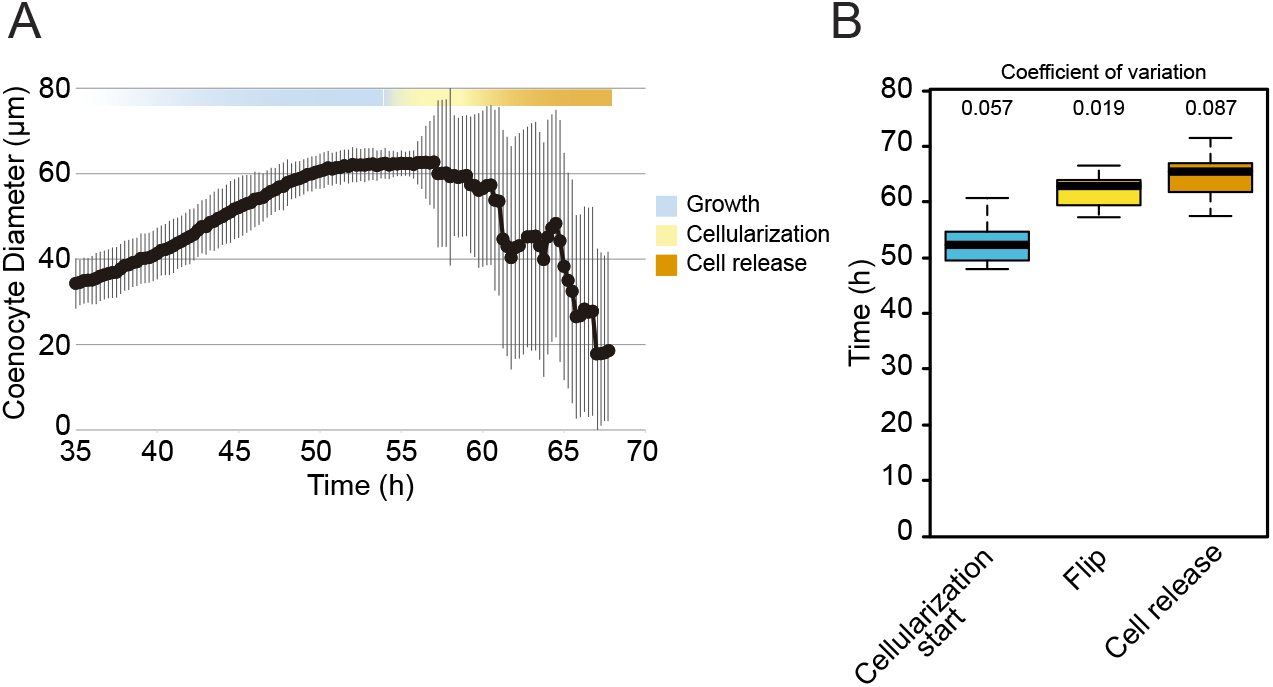
(A) Cell diameter over time of 65 single cell traces aligned to time. Variability increases from 54h onwards due to asynchronous release of new born cells. (B) The average time at which cells start cellularization, undergo flip or release new-born cells in a bulk culture. (N°cells = 100).

**Figure S2.**
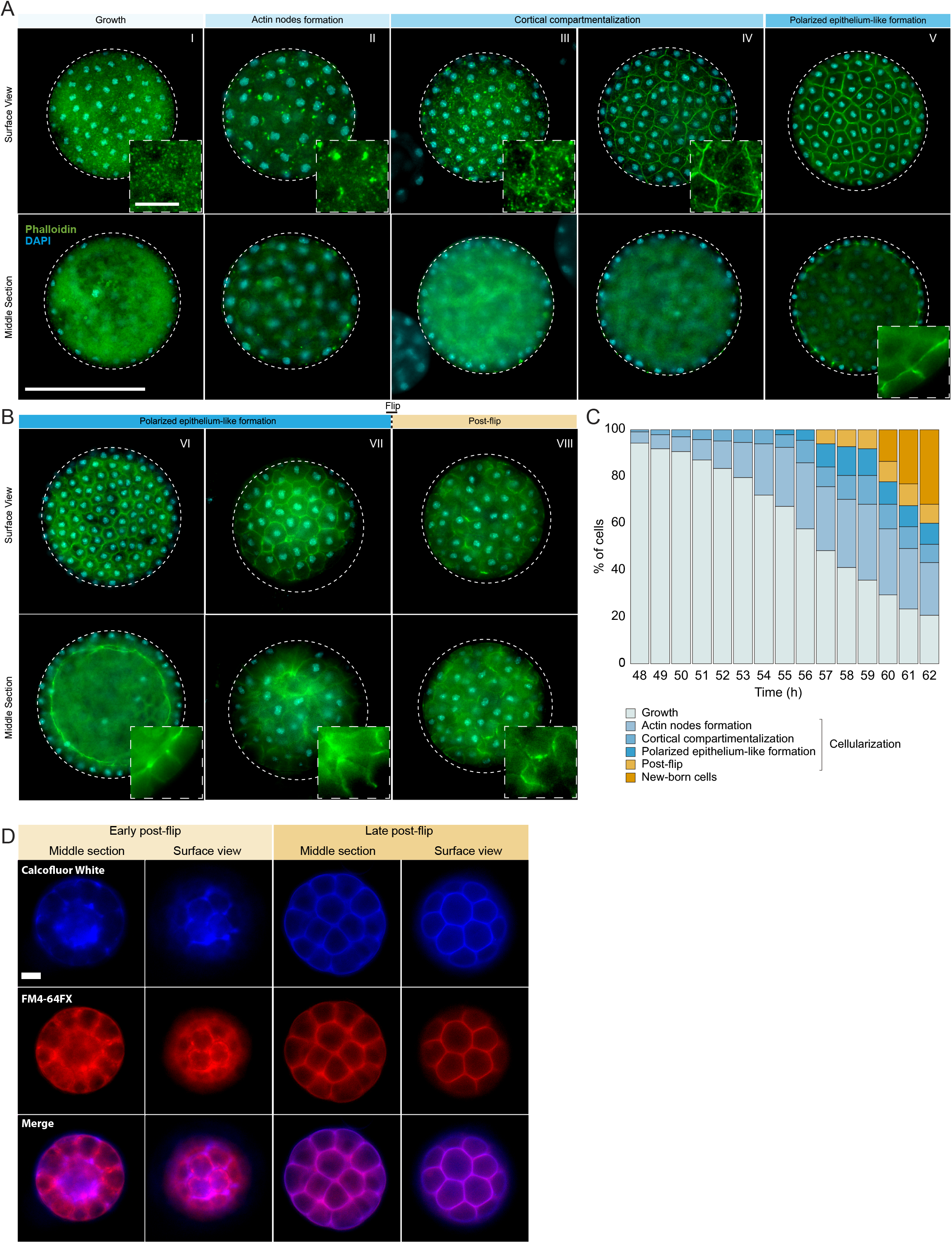
(A) Epifluorescence microscopy images of the cellularization process using phalloidin and DAPI. Arrows show the different actin structures during the different stages of cellularization. Bars, 50μm for whole image and 10μm for the zoom. (B) fraction of cells exhibiting different actin and cellular structures throughout cellularization. The temporal order of actin structures show that the stages occur sequentially. (N°cells = 300) (C) Cell wall formation is observed only post-flip. Cells were fixed and co-stained with the membrane dye FM4-64FX and the cell-wall dye calcofluor white and imaged using an epifluorescence microscope. Bar, 10μm.

**Figure S3.**
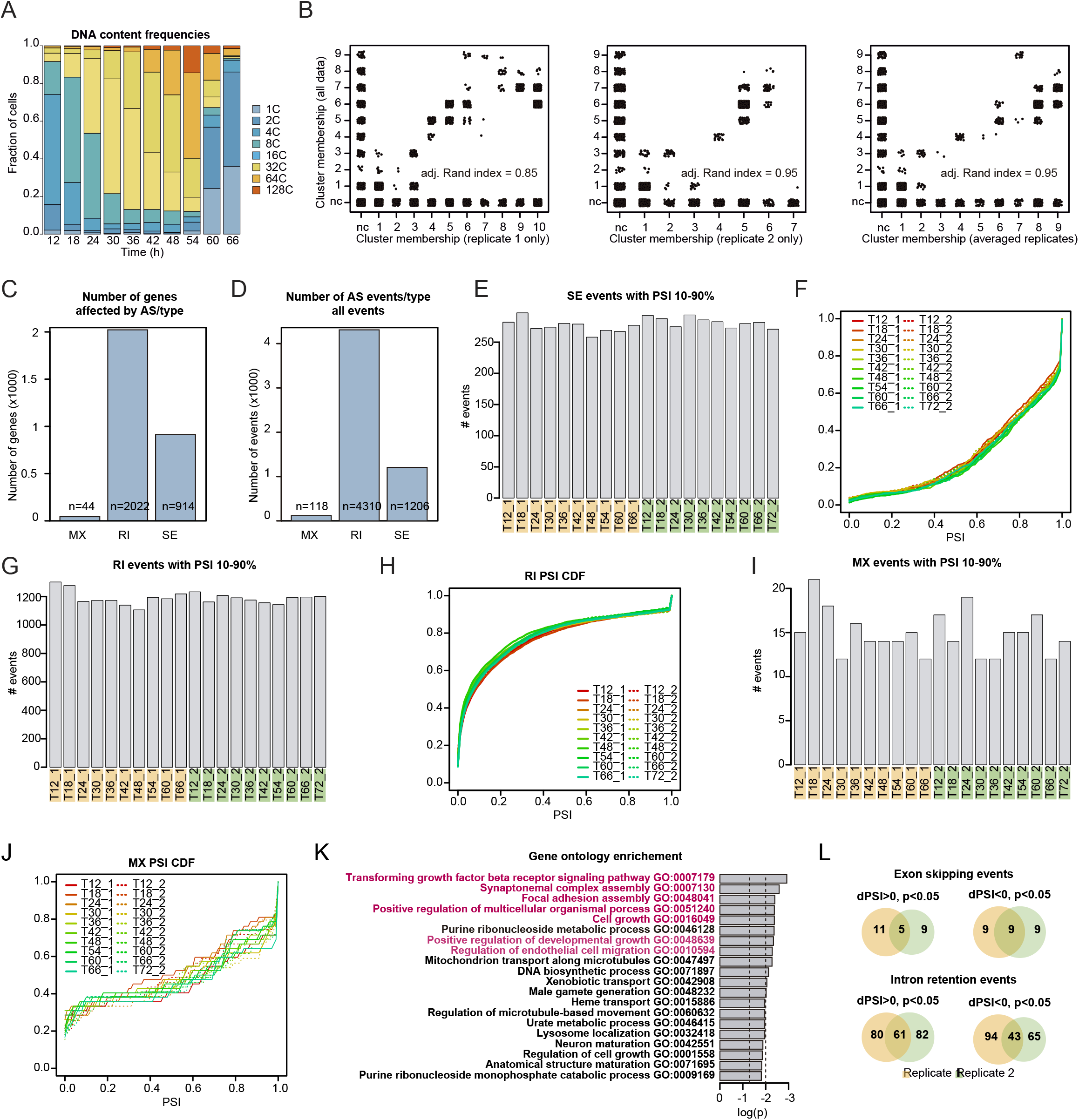
(A) Distributions of nuclear content of *S. arctica* cells across the life cycle in the samples used for transcriptional profiling of a representative replicate, measured by flow cytometry. (B) Cluster membership comparison of transcripts clustering with the entire dataset with samples from either replicate only or averaged abundances of both replicates. The figures and calculated adjusted Rand index demonstrate that cluster memberships are robust. nc - not clustered (genes were not included in either gene expression cluster). (C) number of genes affected by the three types of alternative splicing. MX - mutually exclusive exons; RI - intron retention; SE - exon skipping. (D) number of alternative splicing events. Labels same as in (C) (E-J) Dynamics of alternative splicing across the *S. arctica* life cycle. Number of events per sample and cumulative distribution function (CDF) for exon skipping (SE), intron retention (RI) and mutually exclusive exons (MX). (K) Gene ontology enrichment of genes affected by alternative splicing, calculated by elim method using Bonferroni correction (L) Venn diagram of alternative splicing events that differ significantly between the growth and cellularization stage. dPSI - differential percentage spliced-in.

**Figure S4.**
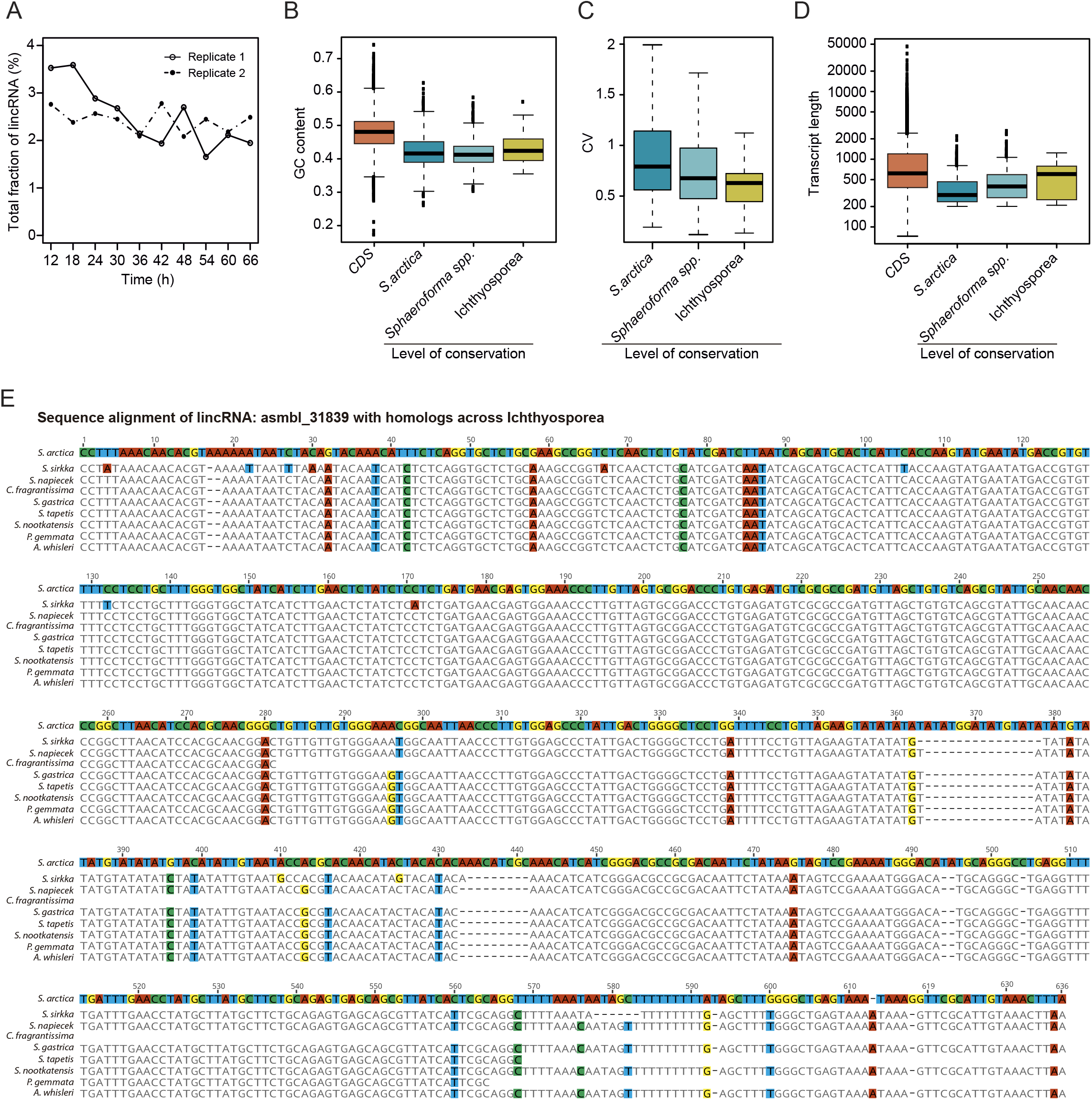
(A) Fraction of total abundance of long intergenic non-coding RNA transcripts (lincRNAs) among all transcripts. (B) Comparison of GC content of lincRNAs binned by degree of conservation with coding genes (CDS). (C) Coefficient of variation of expression across the S. arctica life cycle of lincRNAs binned by degree of conservation. (D) Transcript length of lincRNAs binned by degree of conservation and coding genes. (E) Sequence alignment of lincRNA: asmbl_31839 with homologs across ichthyosporea.

**Figure S5.**
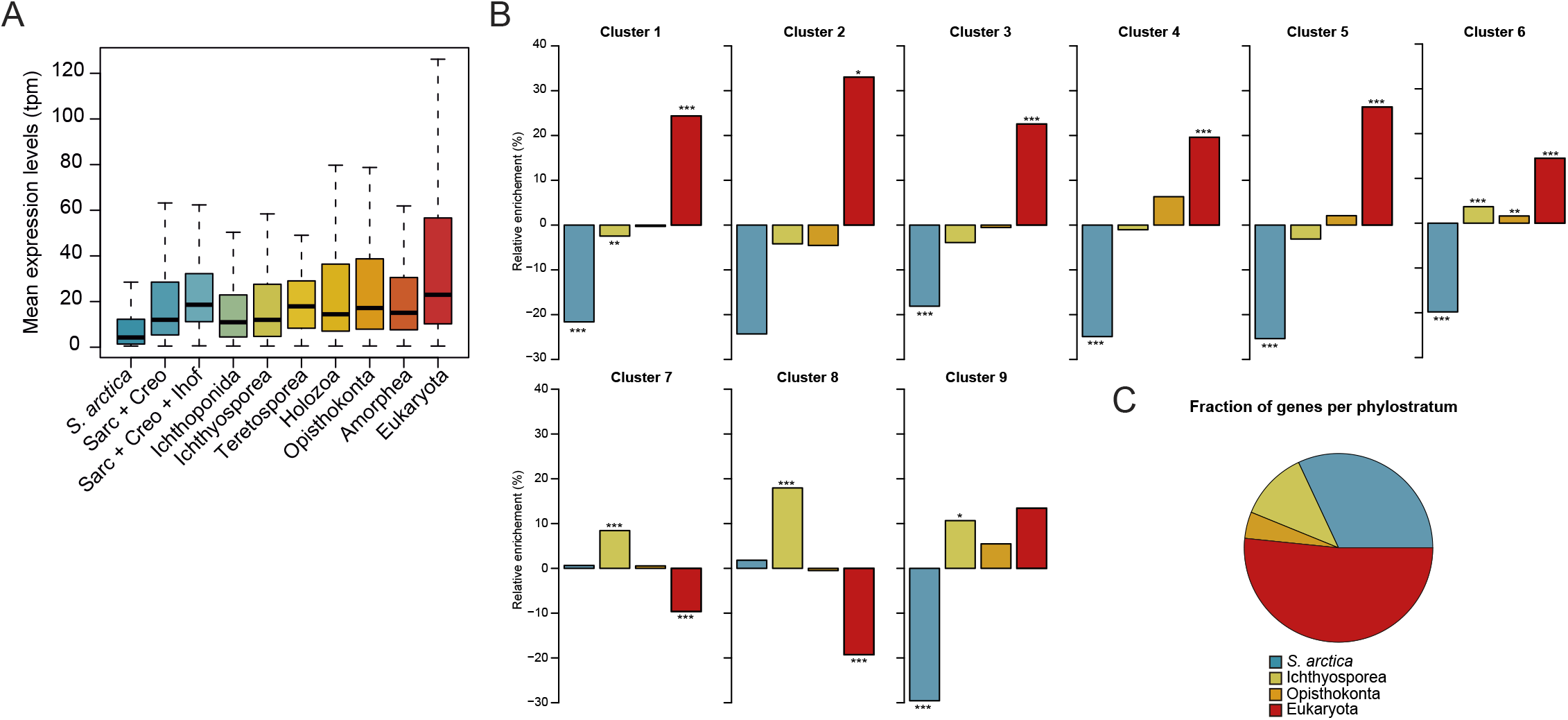
(A) Mean gene expression across the *S. arctica* life cycle, binned by phylostratum. (B) Relative enrichment of gene phylostrata in within gene expression clusters. Original 10 phylostrata were collapsed into 4. (* - p < 0.05, ** - p < 0.01, *** - p < 0.001, Fisher exact test) (C) Fraction of genes per phylostratum used in (B)

**Figure S6.**
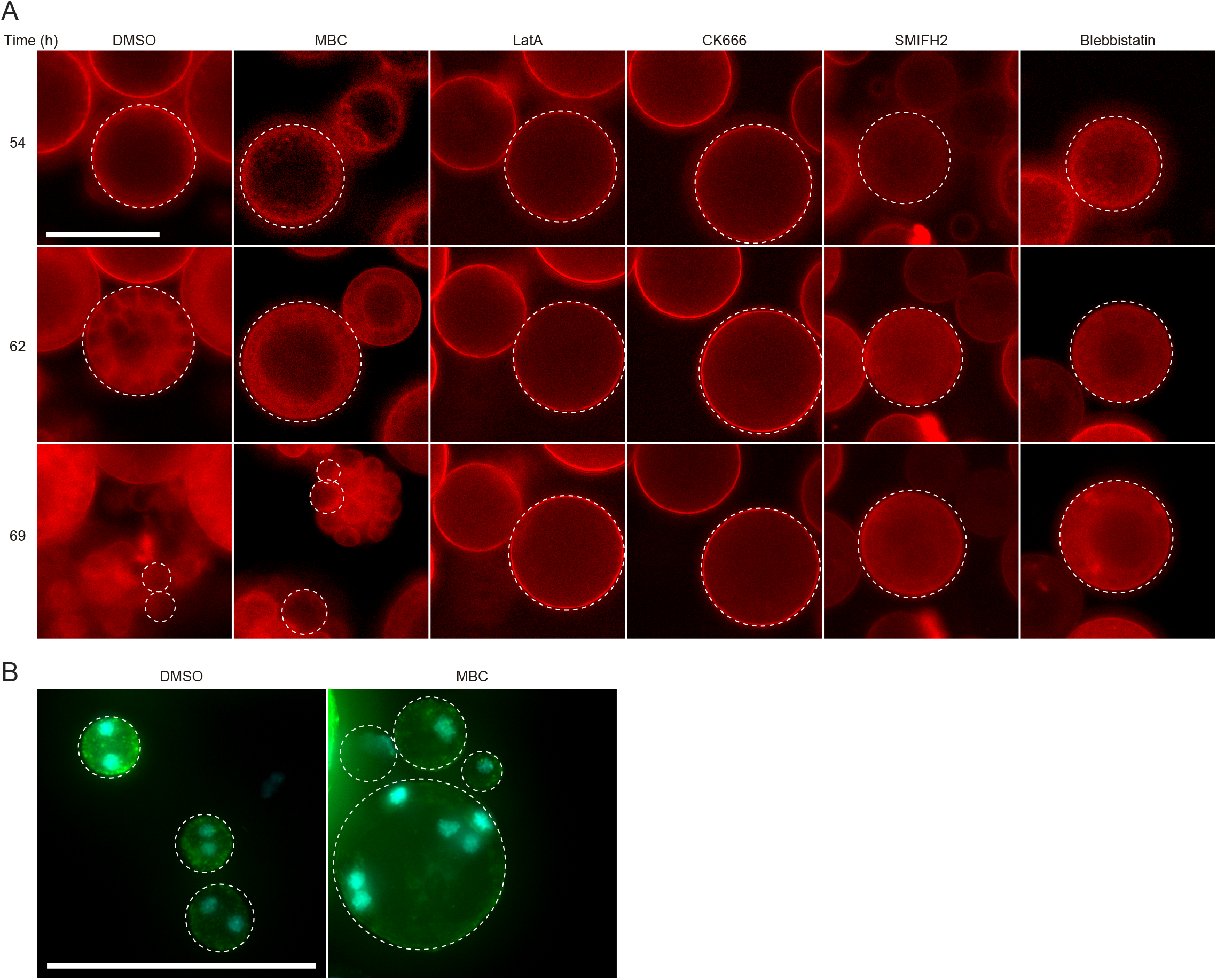
(A) Depolymerization of microtubules and actin, as well as inhibition of Arp2/3, formins and myosin II affect plasma membrane invaginations. Synchronized cells of *S. arctica*, pre-grown for 54 hours, were imaged for 24 hours in presence of multiple pharmacological inhibitors (DMSO as a control, MBC, Latrunculin A, CK666, SMIFH2, Blebbistatin). Note the different delay in cell release and the cell size variability of new born cells after treatment with MBC. Bar, 50μm. (B) Depolymerization of microtubules generates new-born cells with uneven size and number of nuclei. Synchronized cells of *S. arctica*, pre-grown for 54 hours, were imaged after 24 hours in presence MBC or DMSO. Note the different sizes and uneven number of nuclei of MBC-treated new born cells.

**Table S1.** Metrics of the *S. arctica* genome assemblies.

| Metrics | Version 1 (Grau-Bove et al. 2017) | Version 2.1 (this study) |
| --- | --- | --- |
| Total sequence length | 121,588,341 bp | 115,261,641 bp |
| Number of supercontigs | 15,618 | 737 |
| Supercontig N50 | 64,598 bp | 243,943 bp |
| Longest supercontig | 357,797 bp | 776,519 bp |

**Metrics of the *S. arctica* genome assemblies (without the mitochondrial genome)**
| Metrics | Version 1 (Grau-Bove et al. 2017) | Version 2.1 (this study) |
| --- | --- | --- |
| Total sequence length | 121,588,341 bp | 115,079,836 bp |
| Number of supercontigs | 15,618 | 732 |
| Supercontig N50 | 64,598 bp | 244,210 bp |
| Longest supercontig | 357,797 bp | 776,519 bp |

**Table S2.**
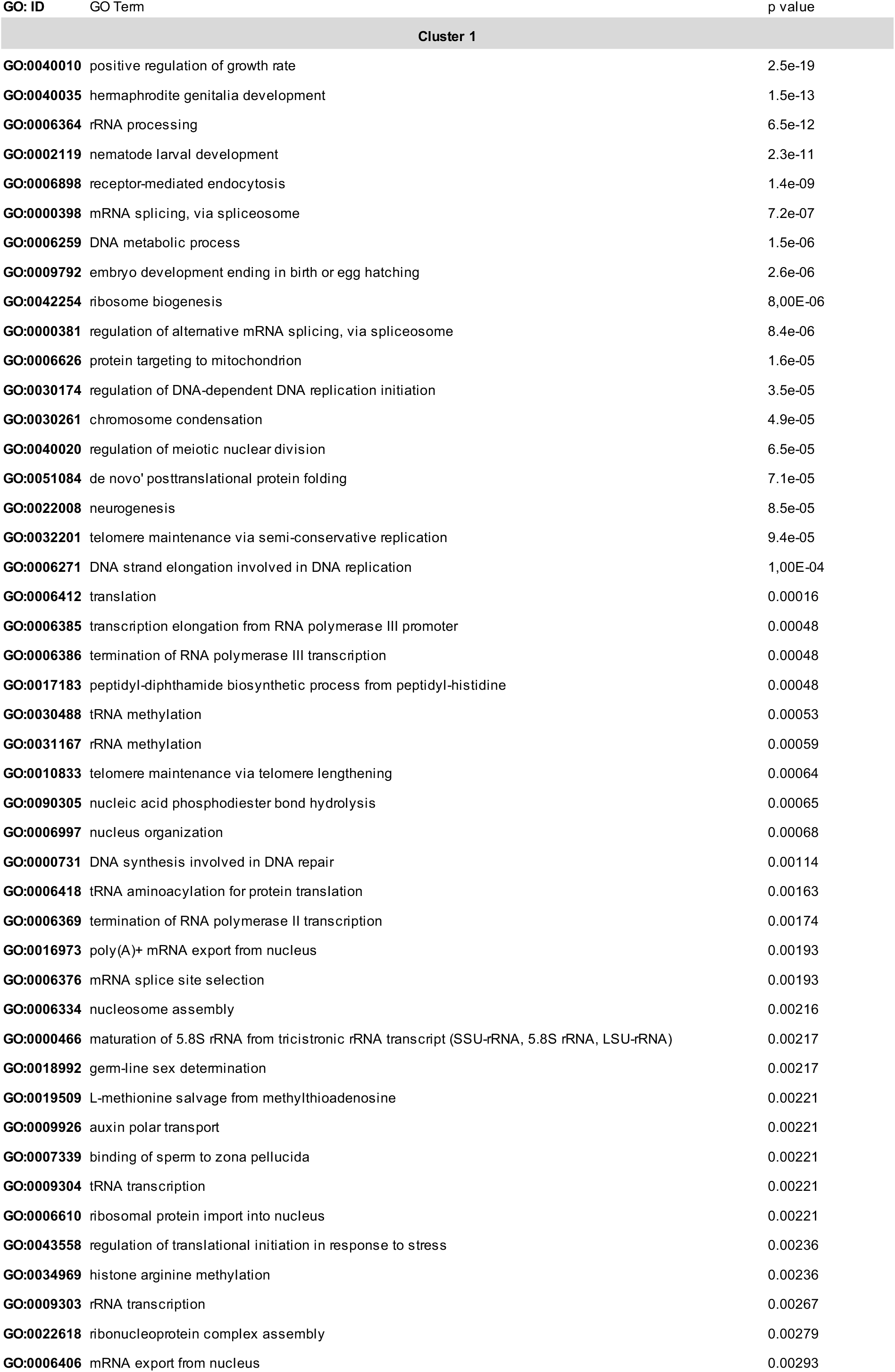
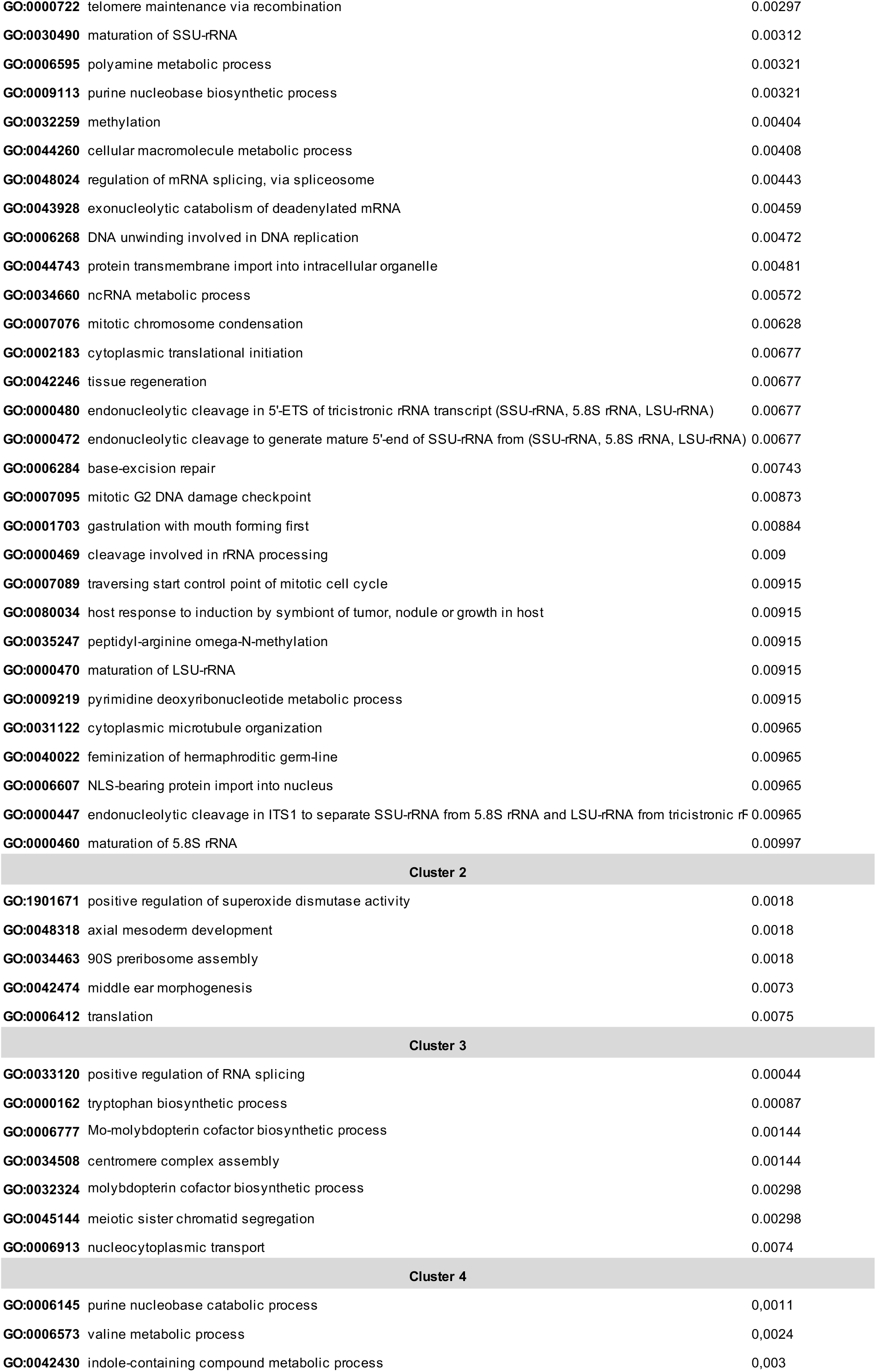
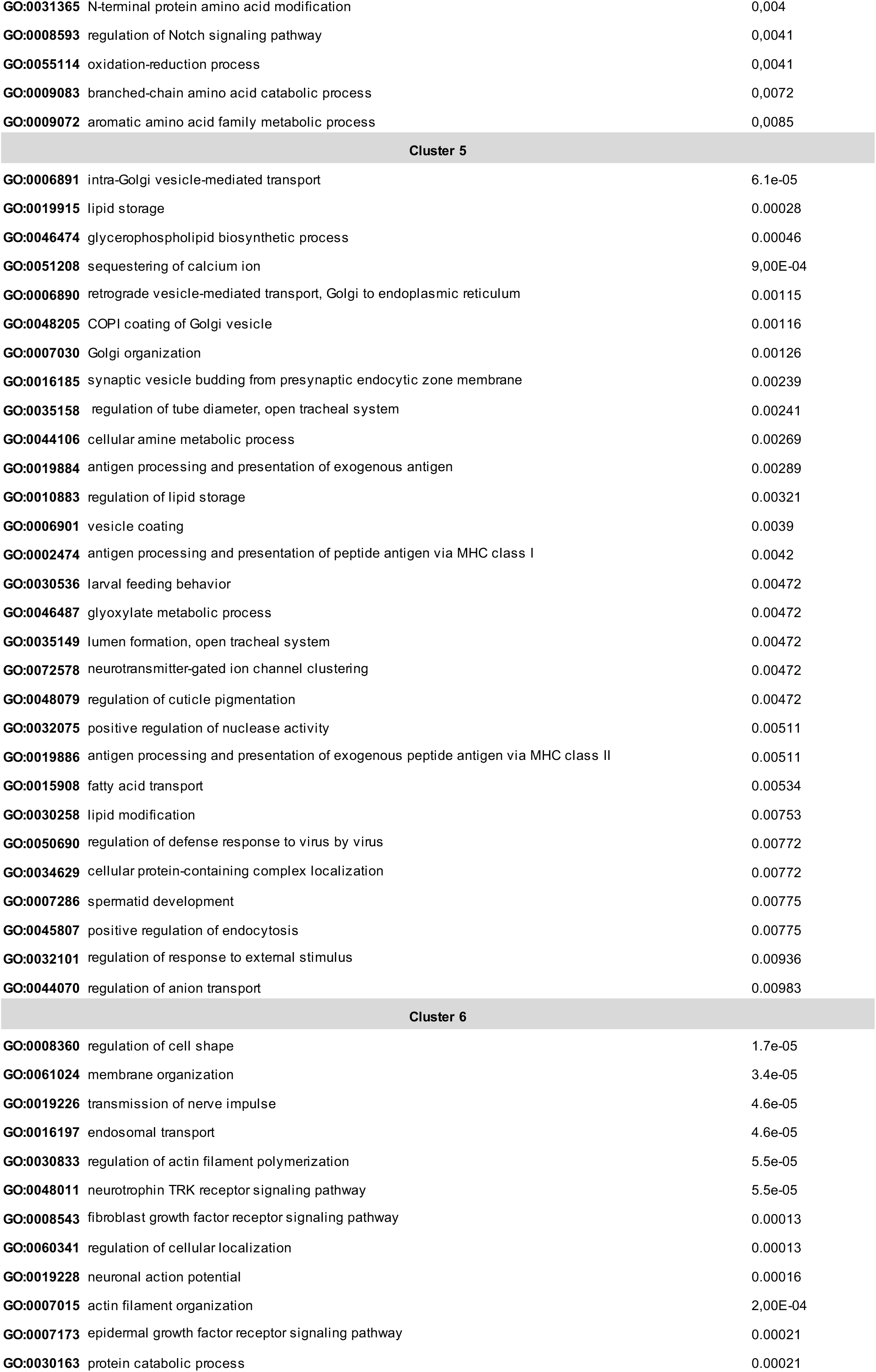
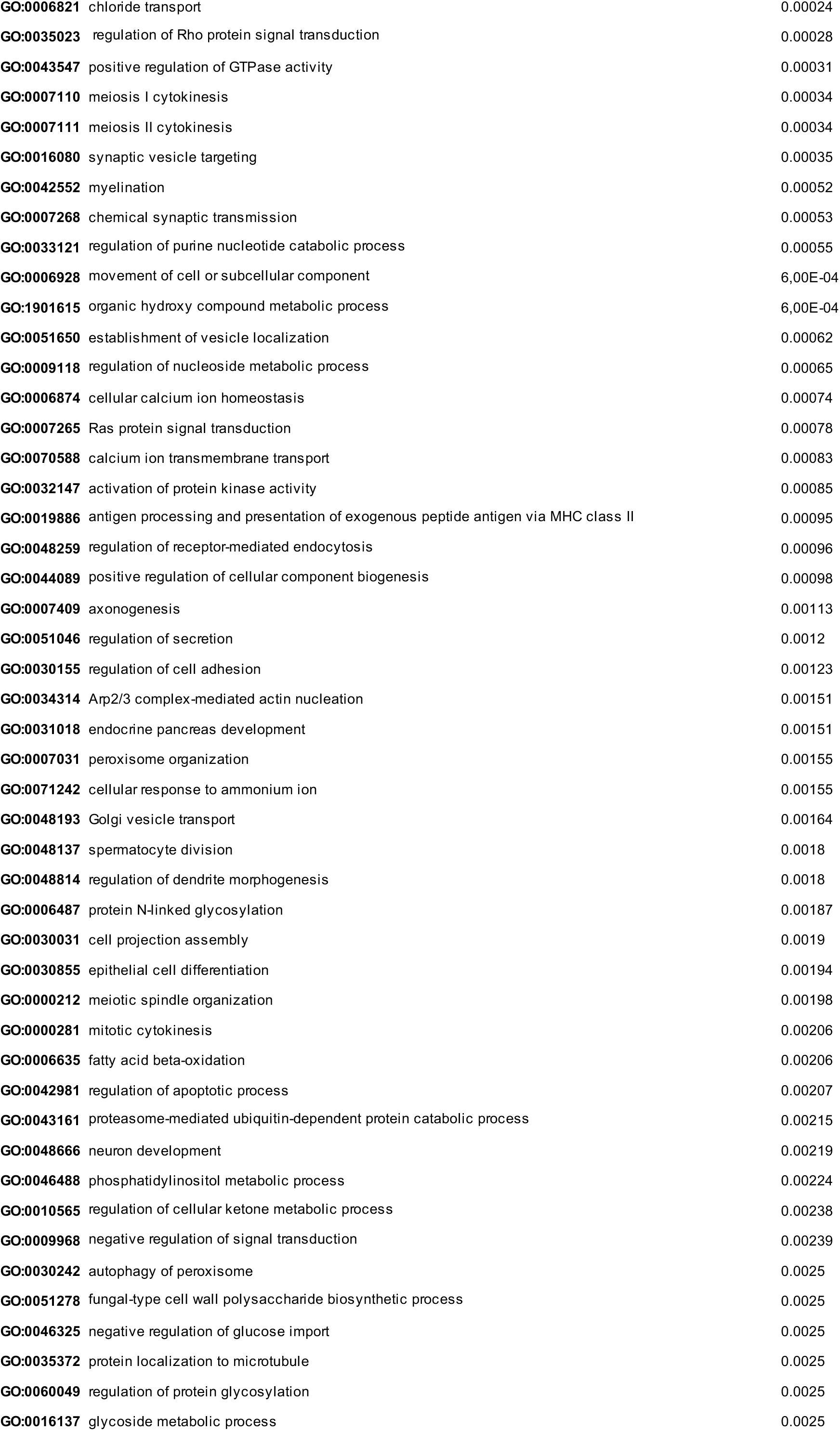
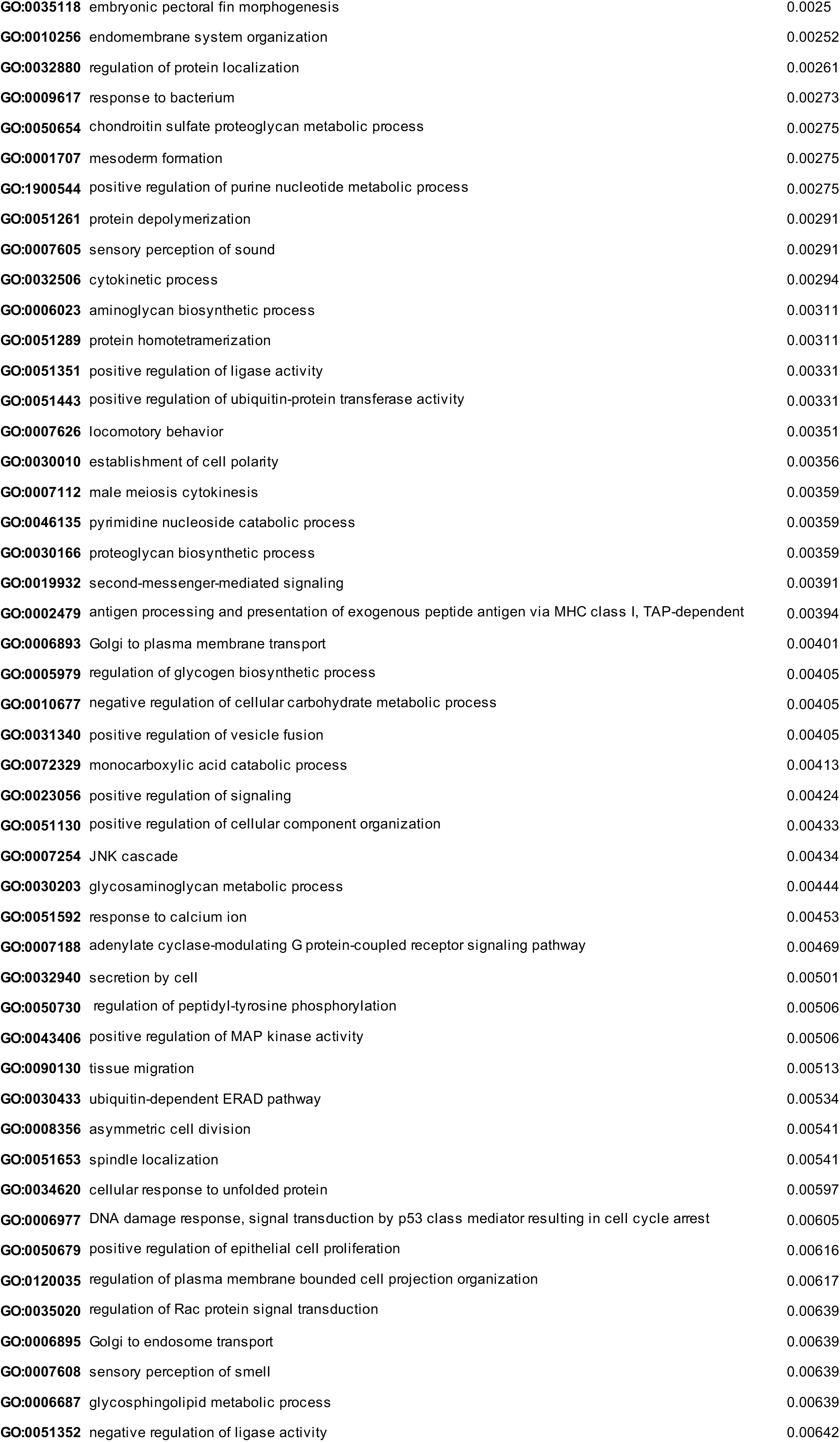
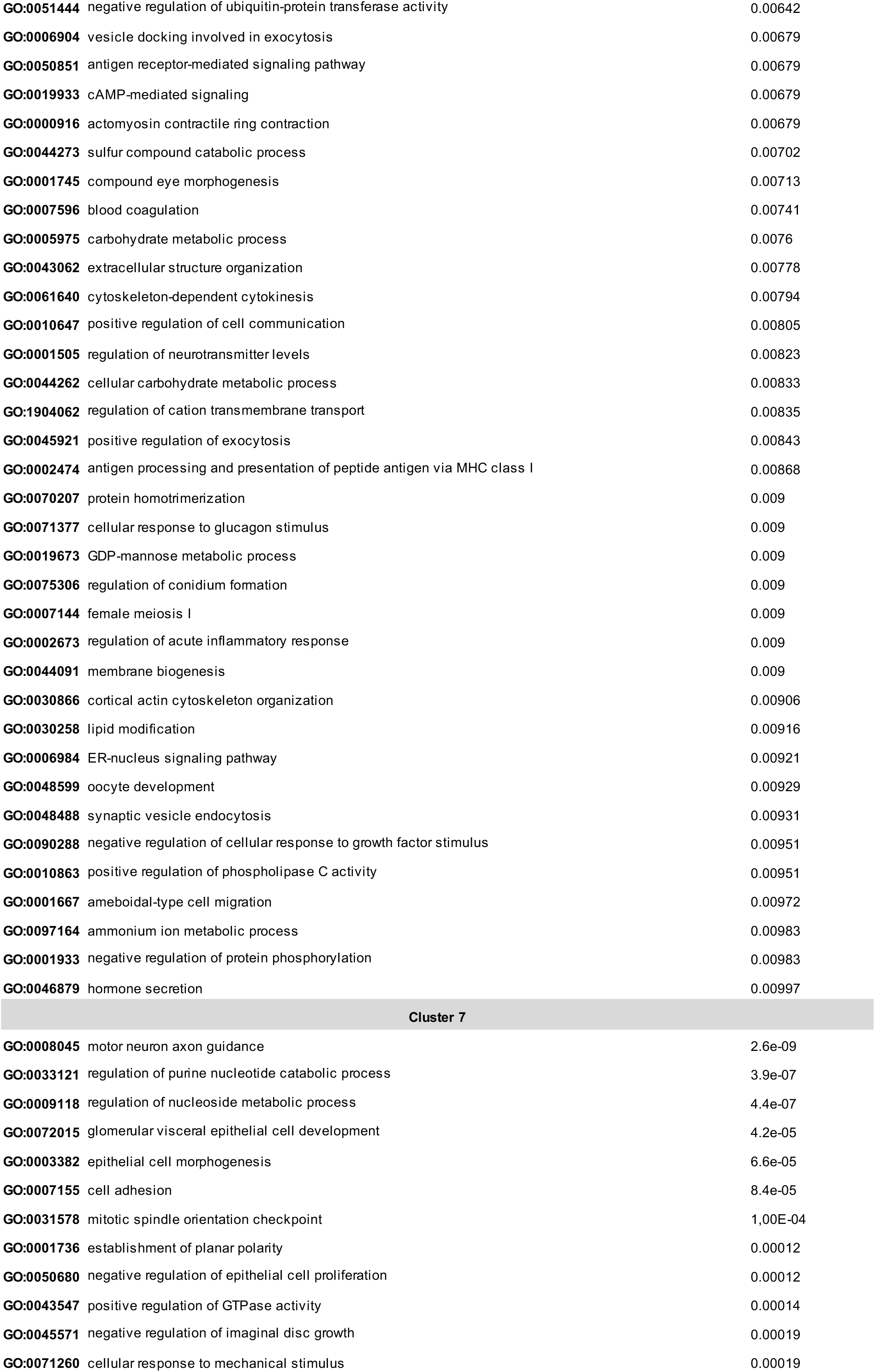
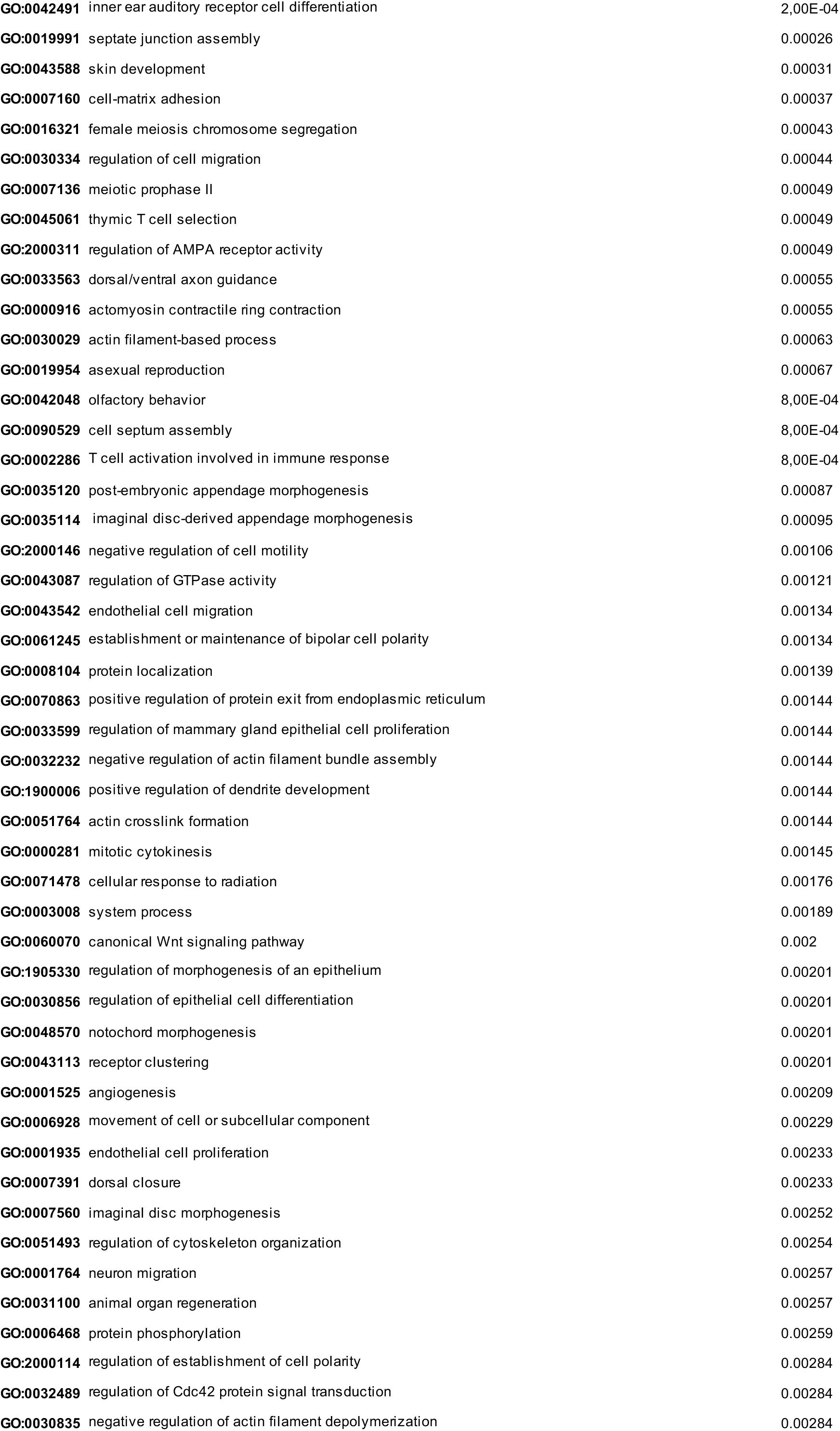
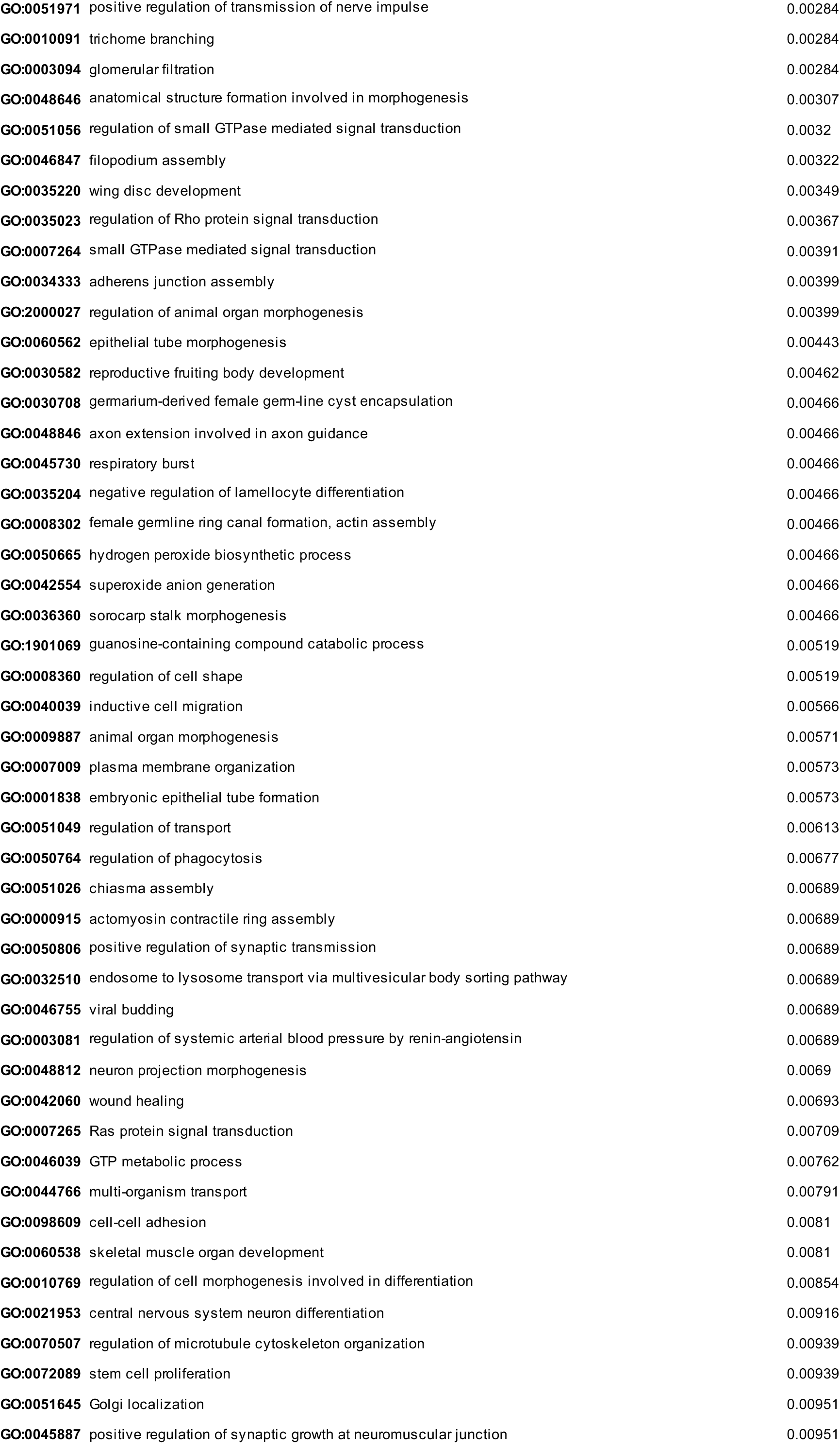
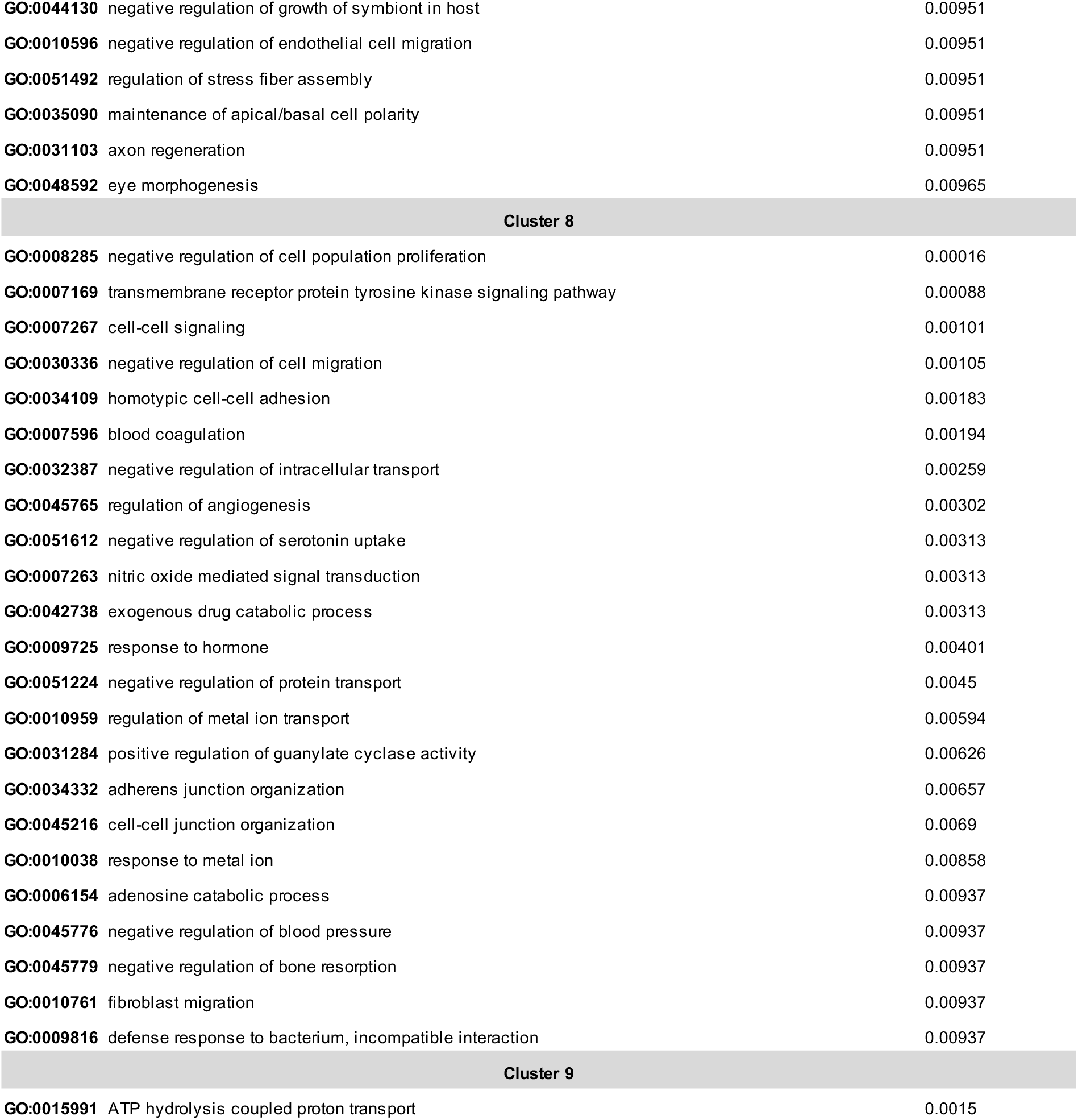
Gene ontology enrichement of expression clusters.

| GO: ID | GO Term | p value |
| --- | --- | --- |
| Cluster 1 |  |  |
| GO:0040010 | positive regulation of growth rate | 2.5e-19 |
| GO:0040035 | hermaphrodite genitalia development | 1.5e-13 |
| GO:0006364 | rRNA processing | 6.5e-12 |
| GO:0002119 | nematode larval development | 2.3e-11 |
| GO:0006898 | receptor-mediated endocytosis | 1.4e-09 |
| GO:0000398 | mRNA splicing, via spliceosome | 7.2e-07 |
| GO:0006259 | DNA metabolic process | 1.5e-06 |
| GO:0009792 | embryo development ending in birth or egg hatching | 2.6e-06 |
| GO:0042254 | ribosome biogenesis | 8,00E-06 |
| GO:0000381 | regulation of alternative mRNA splicing, via spliceosome | 8.4e-06 |
| GO:0006626 | protein targeting to mitochondrion | 1.6e-05 |
| GO:0030174 | regulation of DNA-dependent DNA replication initiation | 3.5e-05 |
| GO:0030261 | chromosome condensation | 4.9e-05 |
| GO:0040020 | regulation of meiotic nuclear division | 6.5e-05 |
| GO:0051084 | de novo' posttranslational protein folding | 7.1e-05 |
| GO:0022008 | neurogenesis | 8.5e-05 |
| GO:0032201 | telomere maintenance via semi-conservative replication | 9.4e-05 |
| GO:0006271 | DNA strand elongation involved in DNA replication | 1,00E-04 |
| GO:0006412 | translation | 0.00016 |
| GO:0006385 | transcription elongation from RNA polymerase III promoter | 0.00048 |
| GO:0006386 | termination of RNA polymerase III transcription | 0.00048 |
| GO:0017183 | peptidyl-diphthamide biosynthetic process from peptidyl-histidine | 0.00048 |
| GO:0030488 | tRNA methylation | 0.00053 |
| GO:0031167 | rRNA methylation | 0.00059 |
| GO:0010833 | telomere maintenance via telomere lengthening | 0.00064 |
| GO:0090305 | nucleic acid phosphodiester bond hydrolysis | 0.00065 |
| GO:0006997 | nucleus organization | 0.00068 |
| GO:0000731 | DNA synthesis involved in DNA repair | 0.00114 |
| GO:0006418 | tRNA aminoacylation for protein translation | 0.00163 |
| GO:0006369 | termination of RNA polymerase II transcription | 0.00174 |
| GO:0016973 | poly(A)+ mRNA export from nucleus | 0.00193 |
| GO:0006376 | mRNA splice site selection | 0.00193 |
| GO:0006334 | nucleosome assembly | 0.00216 |
| GO:0000466 | maturation of 5.8S rRNA from tricistronic rRNA transcript (SSU-rRNA, 5.8S rRNA, LSU-rRNA) | 0.00217 |
| GO:0018992 | germ-line sex determination | 0.00217 |
| GO:0019509 | L-methionine salvage from methylthioadenosine | 0.00221 |
| GO:0009926 | auxin polar transport | 0.00221 |
| GO:0007339 | binding of sperm to zona pellucida | 0.00221 |
| GO:0009304 | tRNA transcription | 0.00221 |
| GO:0006610 | ribosomal protein import into nucleus | 0.00221 |
| GO:0043558 | regulation of translational initiation in response to stress | 0.00236 |
| GO:0034969 | histone arginine methylation | 0.00236 |
| GO:0009303 | rRNA transcription | 0.00267 |
| GO:0022618 | ribonucleoprotein complex assembly | 0.00279 |
| GO:0006406 | mRNA export from nucleus | 0.00293 |
| GO:0000722 | telomere maintenance via recombination | 0.00297 |
| GO:0030490 | maturation of SSU-rRNA | 0.00312 |
| GO:0006595 | polyamine metabolic process | 0.00321 |
| GO:0009113 | purine nucleobase biosynthetic process | 0.00321 |
| GO:0032259 | methylation | 0.00404 |
| GO:0044260 | cellular macromolecule metabolic process | 0.00408 |
| GO:0048024 | regulation of mRNA splicing, via spliceosome | 0.00443 |
| GO:0043928 | exonucleolytic catabolism of deadenylated mRNA | 0.00459 |
| GO:0006268 | DNA unwinding involved in DNA replication | 0.00472 |
| GO:0044743 | protein transmembrane import into intracellular organelle | 0.00481 |
| GO:0034660 | ncRNA metabolic process | 0.00572 |
| GO:0007076 | mitotic chromosome condensation | 0.00628 |
| GO:0002183 | cytoplasmic translational initiation | 0.00677 |
| GO:0042246 | tissue regeneration | 0.00677 |
| GO:0000480 | endonucleolytic cleavage in 5'-ETS of tricistronic rRNA transcript (SSU-rRNA, 5.8S rRNA, LSU-rRNA) | 0.00677 |
| GO:0000472 | endonucleolytic cleavage to generate mature 5'-end of SSU-rRNA from (SSU-rRNA, 5.8S rRNA, LSU-rRNA) | 0.00677 |
| GO:0006284 | base-excision repair | 0.00743 |
| GO:0007095 | mitotic G2 DNA damage checkpoint | 0.00873 |
| GO:0001703 | gastrulation with mouth forming first | 0.00884 |
| GO:0000469 | cleavage involved in rRNA processing | 0.009 |
| GO:0007089 | traversing start control point of mitotic cell cycle | 0.00915 |
| GO:0080034 | host response to induction by symbiont of tumor, nodule or growth in host | 0.00915 |
| GO:0035247 | peptidyl-arginine omega-N-methylation | 0.00915 |
| GO:0000470 | maturation of LSU-rRNA | 0.00915 |
| GO:0009219 | pyrimidine deoxyribonucleotide metabolic process | 0.00915 |
| GO:0031122 | cytoplasmic microtubule organization | 0.00965 |
| GO:0040022 | feminization of hermaphroditic germ-line | 0.00965 |
| GO:0006607 | NLS-bearing protein import into nucleus | 0.00965 |
| GO:0000447 | endonucleolytic cleavage in ITS1 to separate SSU-rRNA from 5.8S rRNA and LSU-rRNA from tricistronic rF | 0.00965 |
| GO:0000460 | maturation of 5.8S rRNA | 0.00997 |
| Cluster 2 |  |  |
| GO:1901671 | positive regulation of superoxide dismutase activity | 0.0018 |
| GO:0048318 | axial mesoderm development | 0.0018 |
| GO:0034463 | 90S preribosome assembly | 0.0018 |
| GO:0042474 | middle ear morphogenesis | 0.0073 |
| GO:0006412 | translation | 0.0075 |
| Cluster 3 |  |  |
| GO:0033120 | positive regulation of RNA splicing | 0.00044 |
| GO:0000162 | tryptophan biosynthetic process | 0.00087 |
| GO:0006777 | Mo-molybdopterin cofactor biosynthetic process | 0.00144 |
| GO:0034508 | centromere complex assembly | 0.00144 |
| GO:0032324 | molybdopterin cofactor biosynthetic process | 0.00298 |
| GO:0045144 | meiotic sister chromatid segregation | 0.00298 |
| GO:0006913 | nucleocytoplasmic transport | 0.0074 |
| Cluster 4 |  |  |
| GO:0006145 | purine nucleobase catabolic process | 0.0011 |
| GO:0006573 | valine metabolic process | 0.0024 |
| GO:0042430 | indole-containing compound metabolic process | 0.003 |
| GO:0031365 | N-terminal protein amino acid modification | 0,004 |
| GO:0008593 | regulation of Notch signaling pathway | 0,0041 |
| GO:0055114 | oxidation-reduction process | 0,0041 |
| GO:0009083 | branched-chain amino acid catabolic process | 0,0072 |
| GO:0009072 | aromatic amino acid family metabolic process | 0,0085 |
| Cluster 5 |  |  |
| GO:0006891 | intra-Golgi vesicle-mediated transport | 6.1e-05 |
| GO:0019915 | lipid storage | 0.00028 |
| GO:0046474 | glycerophospholipid biosynthetic process | 0.00046 |
| GO:0051208 | sequestering of calcium ion | 9,00E-04 |
| GO:0006890 | retrograde vesicle-mediated transport, Golgi to endoplasmic reticulum | 0.00115 |
| GO:0048205 | COPI coating of Golgi vesicle | 0.00116 |
| GO:0007030 | Golgi organization | 0.00126 |
| GO:0016185 | synaptic vesicle budding from presynaptic endocytic zone membrane | 0.00239 |
| GO:0035158 | regulation of tube diameter, open tracheal system | 0.00241 |
| GO:0044106 | cellular amine metabolic process | 0.00269 |
| GO:0019884 | antigen processing and presentation of exogenous antigen | 0.00289 |
| GO:0010883 | regulation of lipid storage | 0.00321 |
| GO:0006901 | vesicle coating | 0.0039 |
| GO:0002474 | antigen processing and presentation of peptide antigen via MHC class I | 0.0042 |
| GO:0030536 | larval feeding behavior | 0.00472 |
| GO:0046487 | glyoxylate metabolic process | 0.00472 |
| GO:0035149 | lumen formation, open tracheal system | 0.00472 |
| GO:0072578 | neurotransmitter-gated ion channel clustering | 0.00472 |
| GO:0048079 | regulation of cuticle pigmentation | 0.00472 |
| GO:0032075 | positive regulation of nuclease activity | 0.00511 |
| GO:0019886 | antigen processing and presentation of exogenous peptide antigen via MHC class II | 0.00511 |
| GO:0015908 | fatty acid transport | 0.00534 |
| GO:0030258 | lipid modification | 0.00753 |
| GO:0050690 | regulation of defense response to virus by virus | 0.00772 |
| GO:0034629 | cellular protein-containing complex localization | 0.00772 |
| GO:0007286 | spermatid development | 0.00775 |
| GO:0045807 | positive regulation of endocytosis | 0.00775 |
| GO:0032101 | regulation of response to external stimulus | 0.00936 |
| GO:0044070 | regulation of anion transport | 0.00983 |
| Cluster 6 |  |  |
| GO:0008360 | regulation of cell shape | 1.7e-05 |
| GO:0061024 | membrane organization | 3.4e-05 |
| GO:0019226 | transmission of nerve impulse | 4.6e-05 |
| GO:0016197 | endosomal transport | 4.6e-05 |
| GO:0030833 | regulation of actin filament polymerization | 5.5e-05 |
| GO:0048011 | neurotrophin TRK receptor signaling pathway | 5.5e-05 |
| GO:0008543 | fibroblast growth factor receptor signaling pathway | 0.00013 |
| GO:0060341 | regulation of cellular localization | 0.00013 |
| GO:0019228 | neuronal action potential | 0.00016 |
| GO:0007015 | actin filament organization | 2,00E-04 |
| GO:0007173 | epidermal growth factor receptor signaling pathway | 0.00021 |
| GO:0030163 | protein catabolic process | 0.00021 |
| **GO:0006821** | chloride transport | 0.00024 |
| **GO:0035023** | regulation of Rho protein signal transduction | 0.00028 |
| **GO:0043547** | positive regulation of GTPase activity | 0.00031 |
| **GO:0007110** | meiosis I cytokinesis | 0.00034 |
| **GO:0007111** | meiosis II cytokinesis | 0.00034 |
| **GO:0016080** | synaptic vesicle targeting | 0.00035 |
| **GO:0042552** | myelination | 0.00052 |
| **GO:0007268** | chemical synaptic transmission | 0.00053 |
| **GO:0033121** | regulation of purine nucleotide catabolic process | 0.00055 |
| **GO:0006928** | movement of cell or subcellular component | 6,00E-04 |
| **GO:1901615** | organic hydroxy compound metabolic process | 6,00E-04 |
| **GO:0051650** | establishment of vesicle localization | 0.00062 |
| **GO:0009118** | regulation of nucleoside metabolic process | 0.00065 |
| **GO:0006874** | cellular calcium ion homeostasis | 0.00074 |
| **GO:0007265** | Ras protein signal transduction | 0.00078 |
| **GO:0070588** | calcium ion transmembrane transport | 0.00083 |
| **GO:0032147** | activation of protein kinase activity | 0.00085 |
| **GO:0019886** | antigen processing and presentation of exogenous peptide antigen via MHC class II | 0.00095 |
| **GO:0048259** | regulation of receptor-mediated endocytosis | 0.00096 |
| **GO:0044089** | positive regulation of cellular component biogenesis | 0.00098 |
| **GO:0007409** | axonogenesis | 0.00113 |
| **GO:0051046** | regulation of secretion | 0.0012 |
| **GO:0030155** | regulation of cell adhesion | 0.00123 |
| **GO:0034314** | Arp2/3 complex-mediated actin nucleation | 0.00151 |
| **GO:0031018** | endocrine pancreas development | 0.00151 |
| **GO:0007031** | peroxisome organization | 0.00155 |
| **GO:0071242** | cellular response to ammonium ion | 0.00155 |
| **GO:0048193** | Golgi vesicle transport | 0.00164 |
| **GO:0048137** | spermatocyte division | 0.0018 |
| **GO:0048814** | regulation of dendrite morphogenesis | 0.0018 |
| **GO:0006487** | protein N-linked glycosylation | 0.00187 |
| **GO:0030031** | cell projection assembly | 0.0019 |
| **GO:0030855** | epithelial cell differentiation | 0.00194 |
| **GO:0000212** | meiotic spindle organization | 0.00198 |
| **GO:0000281** | mitotic cytokinesis | 0.00206 |
| **GO:0006635** | fatty acid beta-oxidation | 0.00206 |
| **GO:0042981** | regulation of apoptotic process | 0.00207 |
| **GO:0043161** | proteasome-mediated ubiquitin-dependent protein catabolic process | 0.00215 |
| **GO:0048666** | neuron development | 0.00219 |
| **GO:0046488** | phosphatidylinositol metabolic process | 0.00224 |
| **GO:0010565** | regulation of cellular ketone metabolic process | 0.00238 |
| **GO:0009968** | negative regulation of signal transduction | 0.00239 |
| **GO:0030242** | autophagy of peroxisome | 0.0025 |
| **GO:0051278** | fungal-type cell wall polysaccharide biosynthetic process | 0.0025 |
| **GO:0046325** | negative regulation of glucose import | 0.0025 |
| **GO:0035372** | protein localization to microtubule | 0.0025 |
| **GO:0060049** | regulation of protein glycosylation | 0.0025 |
| **GO:0016137** | glycoside metabolic process | 0.0025 |
| **GO:0035118** | embryonic pectoral fin morphogenesis | 0.0025 |
| **GO:0010256** | endomembrane system organization | 0.00252 |
| **GO:0032880** | regulation of protein localization | 0.00261 |
| **GO:0009617** | response to bacterium | 0.00273 |
| **GO:0050654** | chondroitin sulfate proteoglycan metabolic process | 0.00275 |
| **GO:0001707** | mesoderm formation | 0.00275 |
| **GO:1900544** | positive regulation of purine nucleotide metabolic process | 0.00275 |
| **GO:0051261** | protein depolymerization | 0.00291 |
| **GO:0007605** | sensory perception of sound | 0.00291 |
| **GO:0032506** | cytokinetic process | 0.00294 |
| **GO:0006023** | aminoglycan biosynthetic process | 0.00311 |
| **GO:0051289** | protein homotetramerization | 0.00311 |
| **GO:0051351** | positive regulation of ligase activity | 0.00331 |
| **GO:0051443** | positive regulation of ubiquitin-protein transferase activity | 0.00331 |
| **GO:0007626** | locomotory behavior | 0.00351 |
| **GO:0030010** | establishment of cell polarity | 0.00356 |
| **GO:0007112** | male meiosis cytokinesis | 0.00359 |
| **GO:0046135** | pyrimidine nucleoside catabolic process | 0.00359 |
| **GO:0030166** | proteoglycan biosynthetic process | 0.00359 |
| **GO:0019932** | second-messenger-mediated signaling | 0.00391 |
| **GO:0002479** | antigen processing and presentation of exogenous peptide antigen via MHC class I, TAP-dependent | 0.00394 |
| **GO:0006893** | Golgi to plasma membrane transport | 0.00401 |
| **GO:0005979** | regulation of glycogen biosynthetic process | 0.00405 |
| **GO:0010677** | negative regulation of cellular carbohydrate metabolic process | 0.00405 |
| **GO:0031340** | positive regulation of vesicle fusion | 0.00405 |
| **GO:0072329** | monocarboxylic acid catabolic process | 0.00413 |
| **GO:0023056** | positive regulation of signaling | 0.00424 |
| **GO:0051130** | positive regulation of cellular component organization | 0.00433 |
| **GO:0007254** | JNK cascade | 0.00434 |
| **GO:0030203** | glycosaminoglycan metabolic process | 0.00444 |
| **GO:0051592** | response to calcium ion | 0.00453 |
| **GO:0007188** | adenylate cyclase-modulating G protein-coupled receptor signaling pathway | 0.00469 |
| **GO:0032940** | secretion by cell | 0.00501 |
| **GO:0050730** | regulation of peptidyl-tyrosine phosphorylation | 0.00506 |
| **GO:0043406** | positive regulation of MAP kinase activity | 0.00506 |
| **GO:0090130** | tissue migration | 0.00513 |
| **GO:0030433** | ubiquitin-dependent ERAD pathway | 0.00534 |
| **GO:0008356** | asymmetric cell division | 0.00541 |
| **GO:0051653** | spindle localization | 0.00541 |
| **GO:0034620** | cellular response to unfolded protein | 0.00597 |
| **GO:0006977** | DNA damage response, signal transduction by p53 class mediator resulting in cell cycle arrest | 0.00605 |
| **GO:0050679** | positive regulation of epithelial cell proliferation | 0.00616 |
| **GO:0120035** | regulation of plasma membrane bounded cell projection organization | 0.00617 |
| **GO:0035020** | regulation of Rac protein signal transduction | 0.00639 |
| **GO:0006895** | Golgi to endosome transport | 0.00639 |
| **GO:0007608** | sensory perception of smell | 0.00639 |
| **GO:0006687** | glycosphingolipid metabolic process | 0.00639 |
| **GO:0051352** | negative regulation of ligase activity | 0.00642 |
| **GO:0051444** | negative regulation of ubiquitin-protein transferase activity | 0.00642 |
| **GO:0006904** | vesicle docking involved in exocytosis | 0.00679 |
| **GO:0050851** | antigen receptor-mediated signaling pathway | 0.00679 |
| **GO:0019933** | cAMP-mediated signaling | 0.00679 |
| **GO:0000916** | actomyosin contractile ring contraction | 0.00679 |
| **GO:0044273** | sulfur compound catabolic process | 0.00702 |
| **GO:0001745** | compound eye morphogenesis | 0.00713 |
| **GO:0007596** | blood coagulation | 0.00741 |
| **GO:0005975** | carbohydrate metabolic process | 0.0076 |
| **GO:0043062** | extracellular structure organization | 0.00778 |
| **GO:0061640** | cytoskeleton-dependent cytokinesis | 0.00794 |
| **GO:0010647** | positive regulation of cell communication | 0.00805 |
| **GO:0001505** | regulation of neurotransmitter levels | 0.00823 |
| **GO:0044262** | cellular carbohydrate metabolic process | 0.00833 |
| **GO:1904062** | regulation of cation transmembrane transport | 0.00835 |
| **GO:0045921** | positive regulation of exocytosis | 0.00843 |
| **GO:0002474** | antigen processing and presentation of peptide antigen via MHC class I | 0.00868 |
| **GO:0070207** | protein homotrimerization | 0.009 |
| **GO:0071377** | cellular response to glucagon stimulus | 0.009 |
| **GO:0019673** | GDP-mannose metabolic process | 0.009 |
| **GO:0075306** | regulation of conidium formation | 0.009 |
| **GO:0007144** | female meiosis I | 0.009 |
| **GO:0002673** | regulation of acute inflammatory response | 0.009 |
| **GO:0044091** | membrane biogenesis | 0.009 |
| **GO:0030866** | cortical actin cytoskeleton organization | 0.00906 |
| **GO:0030258** | lipid modification | 0.00916 |
| **GO:0006984** | ER-nucleus signaling pathway | 0.00921 |
| **GO:0048599** | oocyte development | 0.00929 |
| **GO:0048488** | synaptic vesicle endocytosis | 0.00931 |
| **GO:0090288** | negative regulation of cellular response to growth factor stimulus | 0.00951 |
| **GO:0010863** | positive regulation of phospholipase C activity | 0.00951 |
| **GO:0001667** | ameboidal-type cell migration | 0.00972 |
| **GO:0097164** | ammonium ion metabolic process | 0.00983 |
| **GO:0001933** | negative regulation of protein phosphorylation | 0.00983 |
| **GO:0046879** | hormone secretion | 0.00997 |
| Cluster 7 |  |  |
| **GO:0008045** | motor neuron axon guidance | 2.6e-09 |
| **GO:0033121** | regulation of purine nucleotide catabolic process | 3.9e-07 |
| **GO:0009118** | regulation of nucleoside metabolic process | 4.4e-07 |
| **GO:0072015** | glomerular visceral epithelial cell development | 4.2e-05 |
| **GO:0003382** | epithelial cell morphogenesis | 6.6e-05 |
| **GO:0007155** | cell adhesion | 8.4e-05 |
| **GO:0031578** | mitotic spindle orientation checkpoint | 1,00E-04 |
| **GO:0001736** | establishment of planar polarity | 0.00012 |
| **GO:0050680** | negative regulation of epithelial cell proliferation | 0.00012 |
| **GO:0043547** | positive regulation of GTPase activity | 0.00014 |
| **GO:0045571** | negative regulation of imaginal disc growth | 0.00019 |
| **GO:0071260** | cellular response to mechanical stimulus | 0.00019 |
| **GO:0042491** | inner ear auditory receptor cell differentiation | 2,00E-04 |
| **GO:0019991** | septate junction assembly | 0.00026 |
| **GO:0043588** | skin development | 0.00031 |
| **GO:0007160** | cell-matrix adhesion | 0.00037 |
| **GO:0016321** | female meiosis chromosome segregation | 0.00043 |
| **GO:0030334** | regulation of cell migration | 0.00044 |
| **GO:0007136** | meiotic prophase II | 0.00049 |
| **GO:0045061** | thymic T cell selection | 0.00049 |
| **GO:2000311** | regulation of AMPA receptor activity | 0.00049 |
| **GO:0033563** | dorsal/ventral axon guidance | 0.00055 |
| **GO:0000916** | actomyosin contractile ring contraction | 0.00055 |
| **GO:0030029** | actin filament-based process | 0.00063 |
| **GO:0019954** | asexual reproduction | 0.00067 |
| **GO:0042048** | olfactory behavior | 8,00E-04 |
| **GO:0090529** | cell septum assembly | 8,00E-04 |
| **GO:0002286** | T cell activation involved in immune response | 8,00E-04 |
| **GO:0035120** | post-embryonic appendage morphogenesis | 0.00087 |
| **GO:0035114** | imaginal disc-derived appendage morphogenesis | 0.00095 |
| **GO:2000146** | negative regulation of cell motility | 0.00106 |
| **GO:0043087** | regulation of GTPase activity | 0.00121 |
| **GO:0043542** | endothelial cell migration | 0.00134 |
| **GO:0061245** | establishment or maintenance of bipolar cell polarity | 0.00134 |
| **GO:0008104** | protein localization | 0.00139 |
| **GO:0070863** | positive regulation of protein exit from endoplasmic reticulum | 0.00144 |
| **GO:0033599** | regulation of mammary gland epithelial cell proliferation | 0.00144 |
| **GO:0032232** | negative regulation of actin filament bundle assembly | 0.00144 |
| **GO:1900006** | positive regulation of dendrite development | 0.00144 |
| **GO:0051764** | actin crosslink formation | 0.00144 |
| **GO:0000281** | mitotic cytokinesis | 0.00145 |
| **GO:0071478** | cellular response to radiation | 0.00176 |
| **GO:0003008** | system process | 0.00189 |
| **GO:0060070** | canonical Wnt signaling pathway | 0.002 |
| **GO:1905330** | regulation of morphogenesis of an epithelium | 0.00201 |
| **GO:0030856** | regulation of epithelial cell differentiation | 0.00201 |
| **GO:0048570** | notochord morphogenesis | 0.00201 |
| **GO:0043113** | receptor clustering | 0.00201 |
| **GO:0001525** | angiogenesis | 0.00209 |
| **GO:0006928** | movement of cell or subcellular component | 0.00229 |
| **GO:0001935** | endothelial cell proliferation | 0.00233 |
| **GO:0007391** | dorsal closure | 0.00233 |
| **GO:0007560** | imaginal disc morphogenesis | 0.00252 |
| **GO:0051493** | regulation of cytoskeleton organization | 0.00254 |
| **GO:0001764** | neuron migration | 0.00257 |
| **GO:0031100** | animal organ regeneration | 0.00257 |
| **GO:0006468** | protein phosphorylation | 0.00259 |
| **GO:2000114** | regulation of establishment of cell polarity | 0.00284 |
| **GO:0032489** | regulation of Cdc42 protein signal transduction | 0.00284 |
| **GO:0030835** | negative regulation of actin filament depolymerization | 0.00284 |
| **GO:0051971** | positive regulation of transmission of nerve impulse | 0.00284 |
| **GO:0010091** | trichome branching | 0.00284 |
| **GO:0003094** | glomerular filtration | 0.00284 |
| **GO:0048646** | anatomical structure formation involved in morphogenesis | 0.00307 |
| **GO:0051056** | regulation of small GTPase mediated signal transduction | 0.0032 |
| **GO:0046847** | filopodium assembly | 0.00322 |
| **GO:0035220** | wing disc development | 0.00349 |
| **GO:0035023** | regulation of Rho protein signal transduction | 0.00367 |
| **GO:0007264** | small GTPase mediated signal transduction | 0.00391 |
| **GO:0034333** | adherens junction assembly | 0.00399 |
| **GO:2000027** | regulation of animal organ morphogenesis | 0.00399 |
| **GO:0060562** | epithelial tube morphogenesis | 0.00443 |
| **GO:0030582** | reproductive fruiting body development | 0.00462 |
| **GO:0030708** | germarium-derived female germ-line cyst encapsulation | 0.00466 |
| **GO:0048846** | axon extension involved in axon guidance | 0.00466 |
| **GO:0045730** | respiratory burst | 0.00466 |
| **GO:0035204** | negative regulation of lamellocyte differentiation | 0.00466 |
| **GO:0008302** | female germline ring canal formation, actin assembly | 0.00466 |
| **GO:0050665** | hydrogen peroxide biosynthetic process | 0.00466 |
| **GO:0042554** | superoxide anion generation | 0.00466 |
| **GO:0036360** | sorocarp stalk morphogenesis | 0.00466 |
| **GO:1901069** | guanosine-containing compound catabolic process | 0.00519 |
| **GO:0008360** | regulation of cell shape | 0.00519 |
| **GO:0040039** | inductive cell migration | 0.00566 |
| **GO:0009887** | animal organ morphogenesis | 0.00571 |
| **GO:0007009** | plasma membrane organization | 0.00573 |
| **GO:0001838** | embryonic epithelial tube formation | 0.00573 |
| **GO:0051049** | regulation of transport | 0.00613 |
| **GO:0050764** | regulation of phagocytosis | 0.00677 |
| **GO:0051026** | chiasma assembly | 0.00689 |
| **GO:0000915** | actomyosin contractile ring assembly | 0.00689 |
| **GO:0050806** | positive regulation of synaptic transmission | 0.00689 |
| **GO:0032510** | endosome to lysosome transport via multivesicular body sorting pathway | 0.00689 |
| **GO:0046755** | viral budding | 0.00689 |
| **GO:0003081** | regulation of systemic arterial blood pressure by renin-angiotensin | 0.00689 |
| **GO:0048812** | neuron projection morphogenesis | 0.0069 |
| **GO:0042060** | wound healing | 0.00693 |
| **GO:0007265** | Ras protein signal transduction | 0.00709 |
| **GO:0046039** | GTP metabolic process | 0.00762 |
| **GO:0044766** | multi-organism transport | 0.00791 |
| **GO:0098609** | cell-cell adhesion | 0.0081 |
| **GO:0060538** | skeletal muscle organ development | 0.0081 |
| **GO:0010769** | regulation of cell morphogenesis involved in differentiation | 0.00854 |
| **GO:0021953** | central nervous system neuron differentiation | 0.00916 |
| **GO:0070507** | regulation of microtubule cytoskeleton organization | 0.00939 |
| **GO:0072089** | stem cell proliferation | 0.00939 |
| **GO:0051645** | Golgi localization | 0.00951 |
| **GO:0045887** | positive regulation of synaptic growth at neuromuscular junction | 0.00951 |
| **GO:0044130** | negative regulation of growth of symbiont in host | 0.00951 |
| **GO:0010596** | negative regulation of endothelial cell migration | 0.00951 |
| **GO:0051492** | regulation of stress fiber assembly | 0.00951 |
| **GO:0035090** | maintenance of apical/basal cell polarity | 0.00951 |
| **GO:0031103** | axon regeneration | 0.00951 |
| **GO:0048592** | eye morphogenesis | 0.00965 |
| Cluster 8 |  |  |
| **GO:0008285** | negative regulation of cell population proliferation | 0.00016 |
| **GO:0007169** | transmembrane receptor protein tyrosine kinase signaling pathway | 0.00088 |
| **GO:0007267** | cell-cell signaling | 0.00101 |
| **GO:0030336** | negative regulation of cell migration | 0.00105 |
| **GO:0034109** | homotypic cell-cell adhesion | 0.00183 |
| **GO:0007596** | blood coagulation | 0.00194 |
| **GO:0032387** | negative regulation of intracellular transport | 0.00259 |
| **GO:0045765** | regulation of angiogenesis | 0.00302 |
| **GO:0051612** | negative regulation of serotonin uptake | 0.00313 |
| **GO:0007263** | nitric oxide mediated signal transduction | 0.00313 |
| **GO:0042738** | exogenous drug catabolic process | 0.00313 |
| **GO:0009725** | response to hormone | 0.00401 |
| **GO:0051224** | negative regulation of protein transport | 0.0045 |
| **GO:0010959** | regulation of metal ion transport | 0.00594 |
| **GO:0031284** | positive regulation of guanylate cyclase activity | 0.00626 |
| **GO:0034332** | adherens junction organization | 0.00657 |
| **GO:0045216** | cell-cell junction organization | 0.0069 |
| **GO:0010038** | response to metal ion | 0.00858 |
| **GO:0006154** | adenosine catabolic process | 0.00937 |
| **GO:0045776** | negative regulation of blood pressure | 0.00937 |
| **GO:0045779** | negative regulation of bone resorption | 0.00937 |
| **GO:0010761** | fibroblast migration | 0.00937 |
| **GO:0009816** | defense response to bacterium, incompatible interaction | 0.00937 |
| Cluster 9 |  |  |
| **GO:0015991** | ATP hydrolysis coupled proton transport | 0.0015 |

**Table S3.**
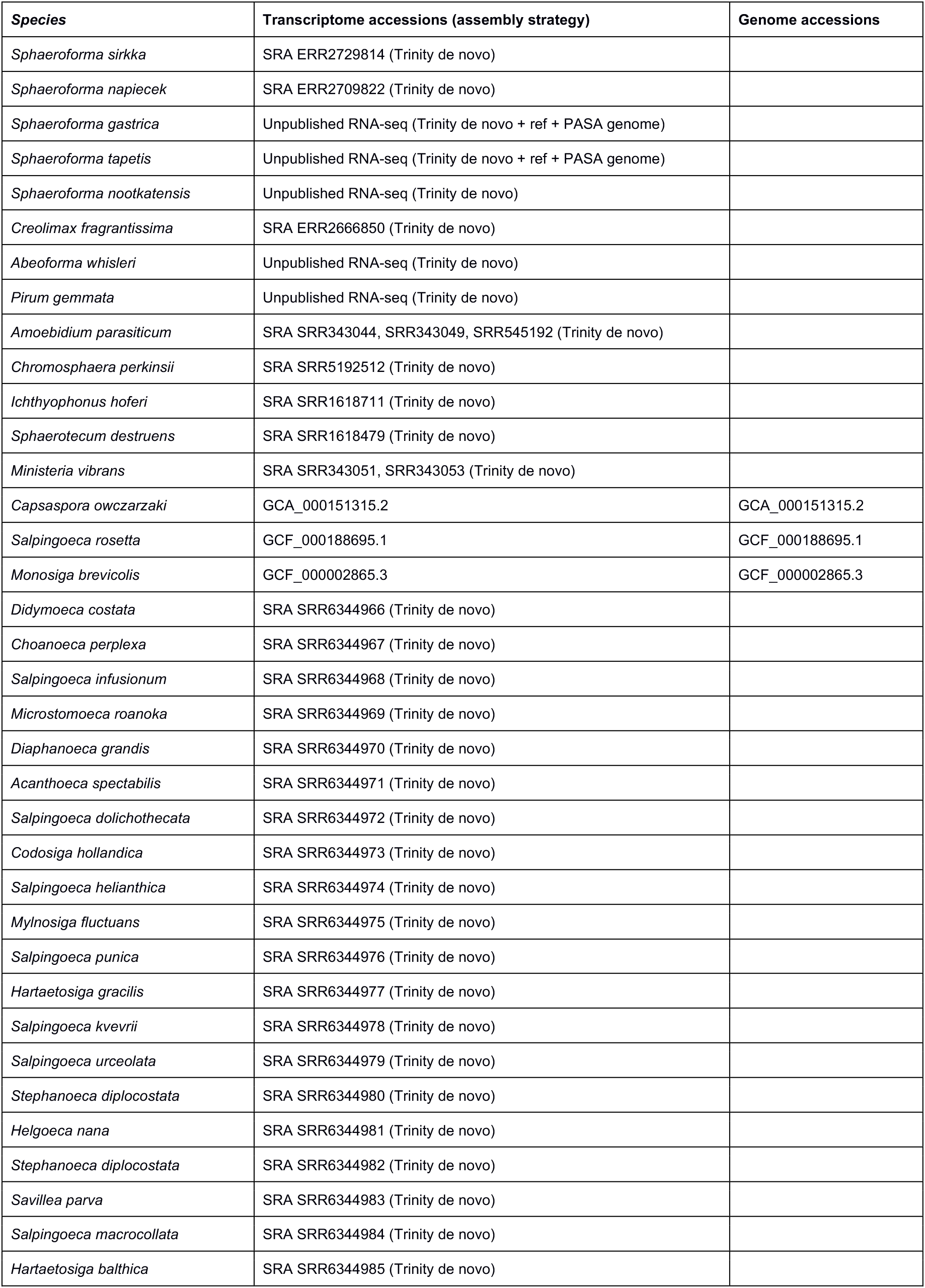
List of genomes and transcriptomes used in the Blastn search to detect conserved S. arctica IncRNAs. SRA indicates the accession number to raw data available from the NCBI Short Read Archive. “Trinity de novo” indicates that the transcriptome was assembled without a reference genome. “Trinity de novo + ref + PASA genome” indicates that the transcriptome was assembled using a reference genome. Assembly details are described in the methods.

## References

Abu-Jamous, B., and Kelly, S. (2018). Clust: automatic extraction of optimal co-expressed gene clusters from gene expression data. Genome Biol. 19, 172.

Adam, J.C., Pringle, J.R., and Peifer, M. (2000). Evidence for Functional Differentiation among Drosophila Septins in Cytokinesis and Cellularization. Mol. Biol. Cell 11, 3123–3135.

Afshar, K., Stuart, B., and Wasserman, S.A. (2000). Functional analysis of the Drosophila diaphanous FH protein in early embryonic development. Development 127, 1887–1897.

Alexa, A., Rahnenführer, J., and Lengauer, T. (2006). Improved scoring of functional groups from gene expression data by decorrelating GO graph structure. Bioinformatics 22, 1600–1607.

Altschul, S.F., Gish, W., Miller, W., Myers, E.W., and Lipman, D.J. (1990). Basic local alignment search tool. J. Mol. Biol. 215, 403–410.

Barter, R.L., and Yu, B. (2018). Superheat: An R Package for Creating Beautiful and Extendable Heatmaps for Visualizing Complex Data. J. Comput. Graph. Stat. 27, 910–922.

Betz, W.J., Mao, F., and Smith, C.B. (1996). Imaging exocytosis and endocytosis. Curr. Opin. Neurobiol. 6, 365–371.

Bolger, A.M., Lohse, M., and Usadel, B. (2014). Trimmomatic: A flexible trimmer for Illumina sequence data. Bioinformatics 30, 2114–2120.

Bonner, J.T. (1998). The origins of multicellularity. Integr. Biol. 1, 27–36.

Braet, F., De Zanger, R., Jans, D., Spector, I., and Wisse, E. (1996). Microfilament-disrupting agent latrunculin A induces and increased number of fenestrae in rat liver sinusoidal endothelial cells: Comparison with cytochalasin B. Hepatology 24, 627–635.

Bråte, J., Adamski, M., Neumann, R.S., Shalchian-Tabrizi, K., and Adamska, M. (2015). Regulatory RNA at the root of animals: dynamic expression of developmental lincRNAs in the calcisponge *Sycon ciliatum.* Proc. R. Soc. B Biol. Sci. 282, 20151746.

Bray, N.L., Pimentel, H., Melsted, P., and Pachter, L. (2016). Near-optimal probabilistic RNA-seq quantification. Nat. Biotechnol. 34, 525–527.

Brunet, T., and King, N. (2017). The Origin of Animal Multicellularity and Cell Differentiation. Dev. Cell 43, 124–140.

Castagnetti, S., Novak, B., and Nurse, P. (2007). Microtubules offset growth site from the cell centre in fission yeast. J. Cell Sci. 120, 2205–2213.

Cooper, J.A., and Kiehart, D.P. (1996). Septins may form a ubiquitous family of cytoskeletal filaments. J. Cell Biol. 134, 1345–1348.

Csuros, M. (2010). Count: evolutionary analysis of phylogenetic profiles with parsimony and likelihood. Bioinformatics 26, 1910–1912.

Dickinson, D.J., Nelson, W.J., and Weis, W.I. (2011). A Polarized Epithelium Organized by predates cadherin and catenin metazoan origins. 1, 1336–1340.

Dickinson, D.J., Nelson, W.J., and Weis, W.I. (2012). An epithelial tissue in Dictyostelium challenges the traditional origin of metazoan multicellularity. BioEssays 34, 833–840.

Dobin, A., Davis, C.A., Schlesinger, F., Drenkow, J., Zaleski, C., Jha, S., Batut, P., Chaisson, M., and Gingeras, T.R. (2012). STAR: ultrafast universal RNA-seq aligner. Bioinformatics 29, 15–21.

Domazet-Lošo, T., and Tautz, D. (2010). A phylogenetically based transcriptome age index mirrors ontogenetic divergence patterns. Nature 468, 815–819.

Eilbeck, K., Lewis, S.E., Mungall, C.J., Yandell, M., Stein, L., Durbin, R., and Ashburner, M. (2005). The Sequence Ontology: a tool for the unification of genome annotations. Genome Biol 6, R44.

Emms, D.M., and Kelly, S. (2015). OrthoFinder: solving fundamental biases in whole genome comparisons dramatically improves orthogroup inference accuracy. Genome Biol. 16, 1–14.

Farrell, J.A., and O’Farrell, P.H. (2014). From Egg to Gastrula: How the Cell Cycle Is Remodeled During the *Drosophila* Mid-Blastula Transition. Annu. Rev. Genet. 48, 269–294.

Gaiti, F., Fernandez-Valverde, S.L., Nakanishi, N., Calcino, A.D., Yanai, I., Tanurdz¡c, M., and Degnan, B.M. (2015). Dynamic and widespread lncRNA expression in a sponge and the origin of animal complexity. Mol. Biol. Evol. 32, 2367–2382.

Giansanti, M.G., Bonaccorsi, S., Williams, B., Williams, E. V., Santolamazza, C., Goldberg, M.L., and Gatti, M. (1998). Cooperative interactions between the central spindle and the contractile ring during Drosophila cytokinesis. Genes Dev. 12, 396–410.

Grabherr, M.G., Haas, B.J., Yassour, M., Levin, J.Z., Thompson, D.A., Amit, I., Adiconis, X., Fan, L., Raychowdhury, R., Zeng, Q., et al. (2011). Full-length transcriptome assembly from RNA-Seq data without a reference genome. Nat. Biotechnol. 29, 644–652.

Grau-bové, X., Ruiz-Trillo, I., and Irimia, M. (2018). Origin of exon skipping-rich transcriptomes in animals driven by evolution of gene architecture. Genome Biology 19, 135.

Grau-Bové, X., Torruella, G., Donachie, S., Suga, H., Leonard, G., Richards, T.A., and Ruiz-Trillo, I. (2017). Dynamics of genomic innovation in the unicellular ancestry of animals. Elife 6, 1–35.

Gunsalus, K.C., Bonaccorsi, S., Williams, E., Verni, F., Gatti, M., and Goldberg, M.L. (1995). Mutations in twinstar, a Drosophila gene encoding a cofilin/ADF homologue, result in defects in centrosome migration and cytokinesis. J. Cell Biol. 131, 1243–1259.

Haas, B.J., Delcher, A.L., Mount, S.M., Wortman, J.R., Smith, R.K., Hannick, L.I., Maiti, R., Ronning, C.M., Rusch, D.B., Town, C.D., et al. (2003). Improving the Arabidopsis genome annotation using maximal transcript alignment assemblies. Nucleic Acids Res. 31, 5654–5666.

Haas, B.J., Papanicolaou, A., Yassour, M., Grabherr, M., Blood, P.D., Bowden, J., Couger, M.B., Eccles, D., Li, B., Lieber, M., et al. (2013). De novo transcript sequence reconstruction from RNA-seq using the Trinity platform for reference generation and analysis. Nat. Protoc. 8, 1494–1512.

Hehenberger, E., Kradolfer, D., and Kohler, C. (2012). Endosperm cellularization defines an important developmental transition for embryo development. Development 139, 2031–2039.

Hetrick, B., Han, M.S., Helgeson, L.A., and Nolen, B.J. (2013). Small molecules CK-666 and CK-869 inhibit actin-related protein 2/3 complex by blocking an activating conformational change. Chem. Biol. 20, 701–712.

Hezroni, H., Koppstein, D., Schwartz, M.G., Avrutin, A., Bartel, D.P., and Ulitsky, I. (2015). Principles of Long Noncoding RNA Evolution Derived from Direct Comparison of Transcriptomes in 17 Species. Cell Rep. 11, 1110–1122.

Hoff, K.J., Lomsadze, A., Stanke, M., and Borodovsky, M. (2018). BRAKER2: Incorporating Protein Homology Information into Gene Prediction with GeneMark-EP and AUGUSTUS. Plant and Animal Genomes XXVI, January 14th 2018.

Huerta-Cepas, J., Forslund, K., Coelho, L.P., Szklarczyk, D., Jensen, L.J., von Mering, C., and Bork, P. (2017). Fast Genome-Wide Functional Annotation through Orthology Assignment by eggNOG-Mapper. Mol. Biol. Evol. 34, 2115–2122.

Hunter, C., and Wieschaus, E. (2000). Regulated expression of nullo is required for the formation of distinct apical and basal adherens junctions in the Drosophila blastoderm. J. Cell Biol. 150, 391–401.

Kalvari, I., Argasinska, J., Quinones-Olvera, N., Nawrocki, E.P., Rivas, E., Eddy, S.R., Bateman, A., Finn, R.D., and Petrov, A.I. (2017). Rfam 13.0: shifting to a genome-centric resource for non-coding RNA families. Nucleic Acids Res. 46, D335–D342.

Kang, Y.-J., Yang, D.-C., Kong, L., Hou, M., Meng, Y.-Q., Wei, L., and Gao, G. (2017). CPC2: a fast and accurate coding potential calculator based on sequence intrinsic features. Nucleic Acids Res. 45, W12–W16.

Keller, O., Kollmar, M., Stanke, M., and Waack, S. (2011). A novel hybrid gene prediction method employing protein multiple sequence alignments. Bioinformatics 27, 757–763.

Kent, W.J. (2002). BLAT - The BLAST-like alignment tool. Genome Res. 12, 656–664.

Kim, D., Langmead, B., and Salzberg, S.L. (2015a). HISAT: A fast spliced aligner with low memory requirements. Nat. Methods 12, 357–360.

Kim, H.C., Jo, Y.J., Kim, N.H., and Namgoong, S. (2015b). Small molecule inhibitor of formin homology 2 domains (SMIFH2) reveals the roles of the formin family of proteins in spindle assembly and asymmetric division in mouse oocytes. PLoS One 10, 1–15.

Kovács, M., Tóth, J., Hetényi, C., Málnási-Csizmadia, A., and Sellers, J.R. (2004). Mechanism of Blebbistatin Inhibition of Myosin II. J. Biol. Chem. 279, 35557–35563.

Lawrence, M., Huber, W., Pagès, H., Aboyoun, P., Carlson, M., Gentleman, R., Morgan, M.T., and Carey, V.J. (2013). Software for Computing and Annotating Genomic Ranges. PLoS Comput. Biol. 9, 1–10.

Leys, S.P., Nichols, S.A., and Adams, E.D.M. (2009). Epithelia and integration in sponges. Integr. Comp. Biol. 49, 167–177.

Lomsadze, A., Ter-Hovhannisyan, V., Chernoff, Y.O., and Borodovsky, M. (2005). Gene identification in novel eukaryotic genomes by self-training algorithm. Nucleic Acids Res. 33, 6494–6506.

Marshall, W.L., and Berbee, M.L. (2011). Facing Unknowns: Living cultures (Pirum gemmata gen. *nov., sp. nov., and Abeoforma whisleri, gen. nov., sp. nov.) from invertebrate digestive tracts represent an undescribed clade within the unicellular opisthokont lineage ichthyosporea (Mesomycetozoea)*. Protist 162, 33–57.

Marshall, W.L., and Berbee, M.L. (2013). Comparative morphology and genealogical delimitation of cryptic species of sympatric isolates of Sphaeroforma (Ichthyosporea, Opisthokonta). Protist 164, 287–311.

Mazumdar, A., and Mazumdar, M. (2002). How one becomes many: Blastoderm cellularization in Drosophila melanogaster. BioEssays 24, 1012–1022.

McLaren, W., Gil, L., Hunt, S.E., Riat, H.S., Ritchie, G.R.S., Thormann, A., Flicek, P., and Cunningham, F. (2016). The Ensembl Variant Effect Predictor. Genome Biol. 17, 1–14.

Mendoza, L., Taylor, J.W., and Ajello, L. (2002). The class mesomycetozoea: a heterogeneous group of microorganisms at the animal-fungal boundary. Annu. Rev. Microbiol. 56, 315–344.

de Mendoza, A., Suga, H., Permanyer, J., Irimia, M., and Ruiz-Trillo, I. (2015). Complex transcriptional regulation and independent evolution of fungal-like traits in a relative of animals. Elife 4, e08904.

Miller, P.W., Clarke, D.N., Weis, W.I., Lowe, C.J., and Nelson, W.J. (2013). The evolutionary origin of epithelial cell-cell adhesion mechanisms. Curr. Top. Membr. 72, 267–311.

Murray, PS., and Zaidel-Bar, R. (2014). Pre-metazoan origins and evolution of the cadherin adhesome. Biol. Open 3, 1183–1195.

Nurk, S., Bankevich, A., Antipov, D., Gurevich, A.A., Korobeynikov, A., Lapidus, A., Prjibelski, A.D., Pyshkin, A., Sirotkin, A., Sirotkin, Y., et al. (2013). Assembling Single-Cell Genomes and Mini-Metagenomes From Chimeric MDA Products. J. Comput. Biol. 20, 714–737.

Ondracka, A., Dudin, O., and Ruiz-Trillo, I. (2018). Decoupling of Nuclear Division Cycles and Cell Size during the Coenocytic Growth of the Ichthyosporean Sphaeroforma arctica. Curr. Biol. 28, 1964–1969.e2.

Parfrey, L.W., Lahr, D.J.G., Knoll, A.H., and Katz, L.A. (2011). Estimating the timing of early eukaryotic diversification with multigene molecular clocks. Proc. Natl. Acad. Sci. 108, 13624–13629.

Patro, R., Duggal, G., Love, M.I., Irizarry, R.A., and Kingsford, C. (2017). Salmon provides fast and bias-aware quantification of transcript expression. Nat. Methods 14, 417–419.

Pignocchi, C., Minns, G.E., Nesi, N., Koumproglou, R., Kitsios, G., Benning, C., Lloyd, C.W., Doonan, J.H., and Hills, M.J. (2009). ENDOSPERM DEFECTIVE1 Is a Novel Microtubule-Associated Protein Essential for Seed Development in Arabidopsis. Plant Cell 21, 90–105.

Quinlan, A.R., and Hall, I.M. (2010). BEDTools: A flexible suite of utilities for comparing genomic features. Bioinformatics 26, 841–842.

Quint, M., Drost, H.G., Gabel, A., Ullrich, K.K., Bönn, M., and Grosse, I. (2012). A transcriptomic hourglass in plant embryogenesis. Nature 490, 98–101.

Richter, D.J., Fozouni, P., Eisen, M.B., and King, N. (2018). Gene family innovation, conservation and loss on the animal stem lineage. Elife 7, 1–43.

Rodriguez-Boulan, E., and Macara, I.G. (2014). Organization and execution of the epithelial polarity programme. Nat. Rev. Mol. Cell Biol. 15, 225–242.

Royou, A., Sullivan, W., and Karess, R. (2002). Cortical recruitment of nonmuscle myosin II in early syncytial Drosophila embryos. J. Cell Biol. 158, 127–137.

Schneider, C.A., Rasband, W.S., and Eliceiri, K.W. (2012). NIH Image to ImageJ: 25 years of image analysis. Nat. Methods 9, 671–675.

Sorensen, M.B. (2002). Cellularisation in the endosperm of Arabidopsis thaliana is coupled to mitosis and shares multiple components with cytokinesis. Development 129, 5567–5576.

Stevenson, V., Hudson, A., Cooley, L., and Theurkauf, W.E. (2002). Arp2/3-dependent psuedocleavage furrow assembly in syncytial Drosophila embryos. Curr. Biol. 12, 705–711.

Suga, H., and Ruiz-Trillo, I. (2013). Development of ichthyosporeans sheds light on the origin of metazoan multicellularity. Dev. Biol. 377, 284–292.

Trincado, J.L., Entizne, J.C., Hysenaj, G., Singh, B., Skalic, M., Elliott, D.J., and Eyras, E. (2018). SUPPA2: Fast, accurate, and uncertainty-aware differential splicing analysis across multiple conditions. Genome Biol. 19, 1–11.

Walker, B.J., Abeel, T., Shea, T., Priest, M., Abouelliel, A., Sakthikumar, S., Cuomo, C.A., Zeng, Q., Wortman, J., Young, S.K., et al. (2014). Pilon: An integrated tool for comprehensive microbial variant detection and genome assembly improvement. PLoS One 9.

Wang, F., Dumstrei, K., Haag, T., and Hartenstein, V. (2004). The role of DE-cadherin during cellularization, germ layer formation and early neurogenesis in the Drosophila embryo. Dev. Biol. 270, 350–363.

Whisler, H.C. (1968). Developmental Control of Amoebidium parasiticurn Amoeba - cyst In tryptone medium In daphnid extract. Dev. Biol. 17, 562–570.

Wu, T.D., and Watanabe, C.K. (2005). GMAP: A genomic mapping and alignment program for mRNA and EST sequences. Bioinformatics 21, 1859–1875.

Xue, W., Li, J.-T., Zhu, Y.-P., Hou, G.-Y., Kong, X.-F., Kuang, Y.-Y., and Sun, X.-W. (2013). L_RNA_scaffolder: scaffolding genomes with transcripts. BMC Genomics 14, 604

